# Primary and secondary antiviral RNAi responses throughout *Varroa destructor* life stages reveal the vertical transmission of viruses

**DOI:** 10.64898/2026.06.11.731771

**Authors:** James Damayo, Rebecca C. McKee, Philip J. Lester, Antoine Felden, Zoe E. Smeele, Alyson Ashe, Emily J. Remnant

**Affiliations:** School of Life and Environmental Sciences, The University of Sydney, Sydney, Australia; Aola Richards Sydney Insect Hub, The University of Sydney, Sydney, Australia; School of Biological Sciences, Victoria University of Wellington, Wellington, New Zealand; Faculty of Biological and Environmental Sciences, University of Helsinki, Helsinki, Finland

**Author notes:** Address correspondence to Emily Remnant.

**Keywords:** RNA interference, vertical transmission, primary siRNA, secondary siRNA, DWV

## Abstract

One of the most devastating threats to global honey bee health is the ectoparasitic mite and viral vector *Varroa destructor*, yet the transmission dynamics of viruses carried by mites are poorly understood. RNA interference (RNAi) is a major antiviral defence mechanism in invertebrates including *Varroa*, where actively replicating viruses are degraded into virus-derived small interfering RNAs (vsiRNAs). Insects typically produce 20–22 nt vsiRNAs with sense and antisense polarity, however established viral infections in *V. destructor* lead to the production of 24-nt antisense vsiRNA fragments, which could indicate the presence of secondary siRNA synthesis. To better understand viral infection and transmission dynamics in *V. destructor*, we conducted small RNA sequencing of male and female mites throughout development, from egg to reproductive stages. Viral community structure was largely driven by developmental stage, with younger and older life stages clustering separately. We identified five viruses that are consistently degraded into antisense 24-nt vsiRNA across all developmental stages, suggesting that these viruses are transmitted vertically and form part of *Varroa*’s core virome. This includes the highly diverse Varroa destructor virus 2 (VDV-2), for which we observe eight distinct VDV-2 strains that simultaneously co-infect individual mites throughout development. In contrast, sense and antisense 23-nt vsiRNA fragments are generated in response to the honey bee pathogen, *Iflavirus aladeformis* (deformed wing virus A, DWV-A) in eggs, but the vsiRNA size profile transitions to 24-nt antisense fragments at later life stages. Our results suggest that once a virus is first acquired by *Varroa*, a primary 23-nt sense and antisense antiviral response precedes the production of secondary 24-nt antisense vsiRNAs as the infection progresses. We confirm this observation using synthetic dsRNA, which show both primary and secondary siRNA processing, revealing how exogenous dsRNA processing occurs in *Varroa*. These results show distinct primary and secondary antiviral RNAi responses across *V. destructor* life stages and demonstrate how vsiRNA profiles can be used to infer virus transmission routes and long-term persistence within vector populations.

## 1 INTRODUCTION

Parasitic mites and ticks represent a critical group of disease drivers across agricultural, veterinary, and human health sectors (Bhat et al., 2025; Brites-Neto et al., 2015). The ectoparasitic mite *Varroa destructor* is presently considered the leading threat to honey bee (*Apis mellifera*) health worldwide (Noël et al., 2020; Rosenkranz et al., 2010; Traynor et al., 2020). *Varroa* mites compromise honey bee health by feeding on haemolymph and fat body tissue, resulting in nutrient loss, the impairment of immune function, and the increased susceptibility to pathogenic viruses (Han et al., 2024; Ramsey et al., 2019; Rosenkranz et al., 2010; Traynor et al., 2020). As well as weakening bee immunity, mites directly transmit pathogenic viruses through vectoring, drastically altering the viral landscape of *A. mellifera*. The arrival of *Varroa* initially drives increases in prevalence and abundance of many common honey bee viruses such as chronic bee paralysis virus, sacbrood virus (SBV; *Iflavirus sacbroodi)*, black queen cell virus (*Triatovirus nigereginacellulae)*, Kashmir bee virus (*Aparavirus kashmirense)*, and Lake Sinai virus 2 (*Sinaivirus secundum*) (Doublet et al., 2024; Lopes et al., 2024; Mondet et al., 2014). However these viruses are eventually superseded by the deformed wing virus (DWV) (Martin et al., 2012; Mondet et al., 2014), a virus that *Varroa* biologically vectors (Damayo et al., 2023) and is considered the main virus associated with colony declines (Wilfert et al., 2016).

*Varroa* harbours a diverse virome of over 20 viral species (Chang et al., 2025; Kim et al., 2026; Lester et al., 2022). Most of these *Varroa-*associated viruses are single-stranded RNA viruses (ssRNA), with the majority having genomes with positive sense polarity (+ssRNA), along with a smaller number of negative sense genomes (-ssRNA) and DNA viruses. The host range of *Varroa*-associated viruses may be restricted to *V. destructor*, including Varroa destructor virus 2 (VDV-2), VDV-3, VDV-5, and VDV-9 (Damayo et al., 2023; Levin et al., 2016, 2019), whereas others are capable of infecting *A. mellifera*. To date, only five viruses have been demonstrated to replicate in both species: Apis rhabdovirus 1 (ARV-1), ARV-2, Bee Macula-like virus, and two DWV genotypes, DWV-A and DWV-B (Chejanovsky et al., 2014; Damayo et al., 2023; Gisder & Genersch, 2020; Norton et al., 2020; Remnant et al., 2017).

*Varroa destructor* is regularly co-infected by multiple viruses, providing ample opportunities for within-vector viral interactions (Eliash et al., 2022). In other vectors, community composition of the virome influences vector competence (Liu et al., 2025; Patterson et al., 2020; H. Wang et al., 2024), altering the efficiency of viral acquisition, transmission, and maintenance. Viral co-infections may also lead to interesting co-evolutionary outcomes, such as viral competition (Carrillo-Tripp et al., 2016) or facilitation (Le Coupanec et al., 2017). In extreme cases, facilitation can manifest as a satellite-helper virus interaction, where sub-viral satellites are completely dependent on helper viruses for replication and/or transmission (Wrzesińska□Krupa & Obrępalska□Stęplowska, 2025). It is crucial to understand how vector-associated viruses are maintained and disseminated throughout populations. Focusing solely on the viruses a vector transmits to its host ignores the important role played by viruses that infect the vector, understating the full complexity of viral transmission dynamics.

Viral maintenance within vector populations mostly relies on two main transmission pathways: vertical and horizontal transmission. Vertical transmission (parent to offspring) can be transovarial, where germline tissues are infected by viruses so that eggs become infected internally, or transovum, where the outer chorion of the eggs are externally contaminated with viral particles that the larva then acquires following eclosion (Y. Chen et al., 2006; Janjoter et al., 2024). In contrast, horizontal transmission occurs intraspecifically between individuals of the same generation via contact or environmental contamination, or interspecifically between a vector and a host (H. Wang et al., 2024). In some instances, arthropod viruses display mixed-modes of transmission which ensure longer-term viral persistence when suitable hosts are scarce or vectoring frequencies decrease (Lequime & Lambrechts, 2014). Evolutionary trade-offs can drive selection for a dominant transmission pathway. Compared to vertical transmission, horizontally transmitted viruses are expected to exhibit higher rates of replication and virulence as their transmission can occur regardless of the survival of their host, while selection for vertically transmitted viruses typically favours lower pathogenicity as fitness depends on the host surviving until it reaches reproduction (Ebert, 2013).

The transmission pathways of a given virus can also be dependent on differences in pathogenicity governed by alternate strains of the same virus. For instance, in *A. mellifera,* the pathogenicity of DWV-A and -B genotypes differs in experimental and field-based studies (McMahon et al., 2016; Natsopoulou et al., 2017; Norton et al., 2020; Tehel et al., 2019). DWV pathogenicity can be further modulated by mode of transmission (vector-driven or direct injection into haemolymph), within-strain sequence variation, or seasonal and vector-density (Ray et al., 2021; Ryabov et al., 2014; Norton et al., 2020; Natsopoulou et al., 2017; Norton et al., 2021).

Distinguishing between vertical, horizontal or vector-driven transmission routes remains a difficult challenge but is crucial for understanding the evolutionary dynamics of pathogens. A robust method to uncover the viral transmission dynamics within *V. destructor* is to identify molecular signals that demonstrate viral replication across life stages. For most invertebrates, including *V. destructor*, RNA interference (RNAi) is one of the main antiviral immune mechanisms, and it provides a reliable indication of active RNA virus replication (Gammon & Mello, 2015; Xu et al., 2021; Zambon et al., 2006). Double stranded RNA (dsRNA) is produced from viral replication intermediates, which are detected by a Dicer-2 endonuclease that cleaves the long dsRNA into shorter, virus derived small-interfering RNA (vsiRNA) fragments ranging from 21-28 nt (MacRae et al., 2007; Siomi & Siomi, 2009; Weber et al., 2006). These siRNAs associate with an Argonaute-2 protein to form the RNA-induced silencing complex, which degrades complementary viral sequences from cognate viral genomes.

The activation of the antiviral siRNA pathway within invertebrate hosts results in the accumulation of vsiRNA fragments that may differ in size depending on the host organism that the viruses are replicating in. Insects generate vsiRNA fragments ranging from 20-22 nt, depending on the host-species (Chejanovsky et al., 2014; Elbashir et al., 2001; Remnant et al., 2017; Santos et al., 2019). Intriguingly, *V. destructor’s* vsiRNA profiles diverge from other acarines. The tick species *Haemaphysalis longicornis* and *Ixodes scapularis* both produce 22 nt vsiRNA profiles with sense and antisense siRNA fragments (Schnettler et al., 2014; Xu et al., 2021), whereas *Varroa* produces a 24 nt vsiRNA profile, with the majority of fragments of antisense polarity (Remnant et al., 2017).

Along with this primary siRNA biogenesis pathway, plants and nematodes have well-characterised secondary siRNA pathways which amplify the antiviral response. Secondary siRNAs are short and antisense to the viral genome, synthesised by RNA-dependent RNA-polymerases (RdRP) (Sijen et al., 2001; X.-B. Wang et al., 2010). RdRPs are host-produced enzymes that are retained in most metazoan lineages (including nematodes and chelicerates), though notably lost from vertebrates and most insects (Lewis et al., 2018; Nganso et al., 2020; Wassenegger & Krczal, 2006; Zong et al., 2009). In ticks and mites, the function of RdRPs in antiviral processes have not been extensively characterised. In the black-legged tick (*Ixodes scapularis*), RdRPs indirectly modulate the expression of viral transcripts by regulating antiviral immune signalling components (Feng et al., 2023), whereas in spider mites (*Tetranychus urticae*), RdRP activity appears limited, contributing minimally to the processing of exogenous RNA (Mondal et al., 2021). Although direct functional evidence is lacking, the vsiRNA profiles produced during *V. destructor’s* antiviral RNAi response show similar characteristics to secondary siRNAs in *C. elegans*, where a large proportion of virus-derived sRNAs are generated antisense to the viral genome (Ashe et al., 2013; Pak & Fire, 2007), suggesting that *Varroa* may also use RdRPs for secondary siRNA synthesis during viral defence.

In this study, we leverage molecular signals of antiviral mediated degradation throughout *V. destructor* development to characterise the vsiRNA biogenesis pathway and to infer viral transmission routes. Viruses that produce vsiRNA profiles that are consistent across all *Varroa* life stages may be vertically transmitted, whereas viruses acquired during an intermediate life stage may indicate horizontal transmission. Here we identify five viruses that are vertically transmitted and are part of the core *V. destructor* virome. Additionally, we observe a switch between two distinct phases of RNAi-mediated antiviral responses to DWV-A across the *Varroa* life stages, suggesting that DWV acquisition also occurs in embryos, though via an alternate mode of vertical transmission. Finally, we validate the distinct primary and secondary siRNA processing phases observed in DWV-A using synthetic dsRNA against target and non-target gene loci to further understand the processing of exogenous dsRNA and antiviral defence in *Varroa*.

## 2 MATERIALS AND METHODS

### 2.1 Sample collection

*Varroa destructor* life stages were collected from capped *Apis mellifera* brood cells from three colonies at the Victoria University of Wellington, New Zealand during March 2022, and February-March 2023. The age and sex of *V. destructor* were determined based on the number of offspring present in the brood cell, the relative sizes and colour of nymphs, and the age of the parasitised pupae (Cameron Jay, 1963; Rosenkranz et al., 2010). Ten life stages of mites were collected: female reproductive phase foundress, female dispersal phase adult, female young adult, female pre-deutochrysalis, female deutonymph, female protonymph, female eggs, male adult, male protonymph, and male eggs. Female dispersal phase adults were previously sequenced from the same apiary in 2022 (Damayo et al., 2023). Pools of three-four mites were collected for smaller life stages (eggs, protonymphs, deutonymphs, and adult males), whilst single mites were collected for larger-sized life stages (pre-deutochrysalises, young adult females, reproductive phase foundresses, and dispersal phase adults), as summarised in Table S1. Previously, these samples were analysed to characterise life stage miRNAome expression (NCBI BioProject: PRJNA1294995, McKee et al., 2026), however we briefly describe the methods used to generate small RNA (sRNA) libraries below.

### 2.2 Extraction of total RNA from *V. destructor* life stages

Total RNA was extracted from whole *V. destructor* with 100 µL of TRI Reagent® (Zymo Research, USA) for single mite and pooled mite samples. Mites were homogenized with a micropestle, in half the total TRI Reagent®, then the remaining reagent was added. Following homogenization, chloroform was added equal to 1/5^th^ of the total TRI Reagent® volume, and the aqueous phase was extracted. The aqueous phase was incubated at −20°C overnight with 100% isopropanol equal to ½ the total TRI Reagent® volume, and 1 µL of glycogen (20 mg/ml). RNA was then precipitated, and the pellets were washed twice with 75% ethanol. Pellets were resuspended in 5.5 µL of nuclease-free water. The quality and concentration of RNA was quantified with a NanoPhotometer NP80 (Implen, Germany) and stored at −80°C.

### 2.3 Small RNA library preparation

Small RNA libraries were generated from 600 ng of total *V. destructor* RNA using the NEBNext Multiplex Small RNA Library Prep Set for Illumina (NEB), using ½ reactions but otherwise following the manufacturers protocol. Amplified cDNA libraries were purified with the Monarch PCR cleanup kit (New England Biolabs, USA), and bands corresponding to sRNA fragments (∼15-35 bp, equating to approximately 140-160 bp fragments inclusive of incorporated adaptors and indices) were size selected on a 6% TBE Acrylamide gel. The resulting libraries were eluted from the gel overnight then precipitated with 3 M NaOAc (pH 5.5), 100% ethanol, and 2 µL of glycogen, at −80°C for 4 hours. Pellets were resuspended in 11 µL of TE buffer and quantified using Qubit Bioanalyzer (ThermoFisher). Libraries were then sent to the Australian Genome Research Facility for sequencing with Illumina NextSeq 2000 (100 bp, single end reads).

### 2.4 Small RNA sequencing data analyses

Raw sequence read quality was checked with FastQC/0.11.8 (Andrews, 2010). Reads were trimmed for adapters and low-quality sequences with Trim Galore/0.6.10 (Krueger, 2016/2023), with a post-trimmed read length minimum of 15 bp, and a Phred quality score cut-off at 20. Ribosomal RNA (rRNA) reads, *V. destructor* genomic reads (Vdes_3.0, Genbank Accession: GCA_002443255.1) and reads mapping to the *A. mellifera* genome (Amel_HAv3.1, Genbank Accession: GCA_003254395.2) were filtered out sequentially using bowtie2/2.2.5 (Langmead & Salzberg, 2012), using the “--very-sensitive” flag, with the minimum seed-length equal to 15 nt. The remaining filtered reads were then mapped to a custom library of viral genomes (Table S2), containing viruses that have been detected in *V. destructor* and *A. mellifera*. We defined sRNA reads aligning to the viral reference library as virus-derived siRNA (vsiRNA). SAM files were converted to BAM files with samtools/1.6 (H. Li et al., 2009) and alignment file summary statistics were extracted with qualimap/2.2.1 (García-Alcalde et al., 2012) to calculate the percentage identity and percentage of reference sequence coverage for each alignment. BAM files were then imported into Geneious Prime/2025.1.2, to visualise coverage for each alignment, then vsiRNA profiles and genomic coverage were plotted in R/2.4.1 with custom scripts.

To extract viral strain information and identify potential viruses that were not present within the custom virus library, contigs were assembled *de novo* from non-host reads with MEGAHIT/1.1.3 (D. Li et al., 2015). A BLASTn search of the assembled contigs against the nucleotide database (NCBI, accessed 10/2024) was conducted, and outputs were filtered to retain hits that were homologous to sequences within the “Viruses”, “Unclassified”, and “N/A” superkingdoms. For contigs that lacked homology to any sequences in the nucleotide database, we conducted a BLASTx search for viral homology at the amino acid level against the non-redundant protein database (NCBI, accessed 10/2024) with DIAMOND/2.0.9 (Buchfink et al., 2015).

### 2.5 Varroa destructor virus 2 strain assembly

In a previous study examining viruses in *V. destructor* from New Zealand (NZ), we found evidence of multiple diverse VDV-2 strains (Lester et al., 2022). To resolve strain diversity to improve vsiRNA mapping, we performed a *de novo* assembly on reads from twenty *V. destructor* transcriptomes [NCBI BioProject: PRJNA820512; (Lester et al., 2022)] that did not map to the *V. destructor* genome using MEGAHIT. Using the ‘map to reference’ function in GeneiousPrime, we identified contigs matching to three available VDV-2 genomes genomes [VDV-2 Israel (NC_040601; Levin et al., 2016); VDV-2 UK (MK795517; Herrero et al., 2019); VDV-2 China (MW590582; G. Chen et al., 2021)] and retained any contigs that were 5 kb or longer. These contigs were generally <90% similar to the available genome sequences. We then used these contigs as bait to probe for additional shorter matching contigs from within the NZ samples. The process of multiple iterative mapping steps gradually pieced together longer contigs by identifying new overlapping contigs that extended the 5’ and 3’ ends of the original contig. This process uncovered eight distinct VDV-2 variants, which we denoted as VDV-2-alpha, -beta, -gamma, -delta, -epsilon, -zeta, -eta and -theta (Table S3). Finally, to confirm that each variant was present at sufficient loads in individual samples, raw sequencing reads from eight of the NZ *V. destructor* transcriptome samples were mapped to each extended NZ VDV-2 contig using bowtie2.

RNA sequencing data from eight additional *V. destructor* life stage samples from the United States of America (US) was downloaded from NCBI to validate the presence of multiple VDV-2 strains in mites from other locations (NCBI BioProject PRJNA380433; SRA accessions SRR5377263-SRR537770). Reads from each sample were assembled *de novo* using MEGAHIT, and the resulting contigs from all samples were mapped to the NZ VDV-2 variants using the ‘map to reference’ function in GeneiousPrime, followed by generation of consensus sequences to generate a US equivalent for each NZ variant.

We performed a multiple sequence alignment with the available *V. destructor* VDV-2 genomes (Israel; UK, and China), two related viral genomes sequenced from *Varroa jacobsoni* in Papua New Guinea [VJV-2-PNG1 (MT482466); VJV-2-PNG2 (MT482467)] and the eight variants assembled from NZ and the US using MUSCLE (Edgar, 2004), implemented in GeneiousPrime. As some consensus sequences were incomplete, we trimmed the alignment to retain a 4 kb segment of the VDV-2 genome which had coverage from all eight strains in both locations. A maximum-likelihood phylogenetic tree was inferred using IQ-TREE (Nguyen et al., 2015), with the best-fit substitution model of TIM2+F+I+G4 determined by ModelFinder (Kalyaanamoorthy et al., 2017). Branch supports were estimated using ultrafast bootstrap approximation (UFBoot2; Hoang et al., 2018) and SH-aLRT (Guindon et al., 2010).

The NZ and US variant consensus sequences have been submitted to NCBI (Genbank accession numbers PZ380877-PZ380884 and BK083173-BK083180; Table S3).

### 2.6 Viral community composition

To visualise the viral community structure among *V. destructor* life stages, we first normalised vsiRNA reads to reads per million (RPM) of vsiRNA reads. RPM-normalised vsiRNA values were centred log-ratio (CLR) transformed for Principle Component Analysis (PCA) with the “compositions” R package (Boogaart et al., 2025). PERMANOVAs were conducted from a sample vs virus Bray-Curtis dissimilarity matrix, generated with the “vegan” R package (Oksanen et al., 2026), to assess the effects of life stage and sex on the viral community composition. A two-way PERMANOVA was conducted to assess the combined effect of age and sex, each with 999 permutations.

Pairwise correlations between viruses were calculated with Spearman’s rank correlation on CLR-transformed RPM values to account for the compositional structure of data within individual samples. P-values were adjusted using the Benjamini-Hochberg method. The resulting correlation matrix was visualised using a heatmap plotted with the “corrplot” R package (Wei et al., 2024). To visualise the vsiRNA levels across samples, log10-transformed, RPM-normalised vsiRNA values were used for hierarchical clustering using Euclidean distances and complete linkage in “pheatmap” (Kolde, 2025).

### 2.7 Assessing siRNA profiles by antisense indices

We hypothesised that in *Varroa*, generation of a primary antiviral siRNA response generates 23 nt fragments of both sense and antisense direction, and the secondary response results in 24 nt antisense fragments. To determine whether a primary or secondary response is predominant for each virus and at different life stages, we devised an index that calculates the ratio of antisense fragment sizes. By focusing on the antisense fragment size only, we avoid the confounding, non-specific degradation products that we observe in viruses that have a positive-sense RNA genome (Damayo et al., 2023). We calculated the antisense index with the equation below:

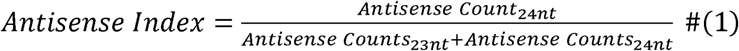

Antisense indices were only calculated for samples in which the combined 23nt and 24nt antisense counts exceeded 50, to avoid distortion of the index due to low read-depth, which resulted in exclusion of 11 samples from DWV-A antisense index analyses.

### 2.8 Quantitative PCR

In all life stage samples, we performed qPCR to quantify the relative load of DWV-A normalised to two reference genes, 18S and NADHD (Campbell et al., 2016). 100 ng of extracted RNA was first treated with ezDNase (Thermo Fisher Scientific), and cDNA was synthesised with the First Strand SuperScript III Reverse Transcriptase kit (Thermo Fisher Scientific) using ½ reactions of the manufacturers protocol, and 25 ng/µl of random hexamer primers. All reactions were conducted in 10 µl PowerUp SYBR Green Master Mix (Applied Biosystems) reactions with 3 ng of template cDNA per well, in triplicate within 96-well plates on a QuantStudio 3 Real-Time PCR System (Applied Biosystems) with the following steps, 50 °C (2 min), 95 °C (2 min), followed by 40 cycles of 95 °C (1 s), 60 °C (30 s), then a melt-curve analysis from 60 °C to 95 °C.

We assessed the primer efficiencies from a standard curve consisting of six 3-fold serial dilutions. Primer efficiencies for this assay ranged from 98%-105% (Table S4). Relative expression of DWV-A was calculated as normalised relative quantities (NRQ) from efficiency-corrected normalisation to the geometric mean of two references genes, with the following equation derived from Hellemans et al. (2007):

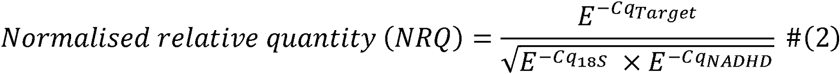

### 2.9 Synthetic dsRNA treatments

To determine how dsRNA is processed by *V. destructor* RNAi machinery, we treated mites with two 800 bp synthetic dsRNA constructs: a targeting dsRNA designed to knock down a Ago2 homologue, and a GFP-dsRNA off-target control (Table S5). Ago2-dsRNA was designed to encompass an exon-exon boundary in the coding sequence of LOC111255253. Both dsRNA constructs were produced via *in vitro* transcription and magnetic bead purification by RNA Greentech LLC (Texas, USA). RNA was concentrated to 5 µg/µl with the Concentrator 5301 (Eppendorf) at 30°C.

To deliver our dsRNA to *V. destructor,* we adapted a protocol first developed by Campbell et al. (2010) for dsRNA-mediated gene knockdown. Foundress mites were collected from capped brood cells from an untreated hive at the Victoria University of Wellington, New Zealand, in February 2024. For each treatment, twelve mites were immersed in three tubes containing 20 µl of 2.5 µg/µl dsRNA in 0.9% NaCl or only 0.9% NaCl for a no-dsRNA control. To ensure that mites were fully submerged, tubes were briefly spun with a mini centrifuge then incubated overnight (∼14 hours) at 4°C. Mites were then allowed to recover at ∼30°C, >75% RH in petri dishes for 3 hours then transferred to gelatine capsules (Size 00, HerbalTech) containing white-eyed worker pupae for 48 hours. Mites that responded to gentle agitation with a fine paintbrush were defined as alive, then stored at −80°C.

Total RNA was extracted from all mites individually following the methods described above, and qPCR was used to validate the success of dsRNA knockdown of *Ago2* (LOC111255253). Relative expression was calculated with the Pfaffl method (Pfaffl, 2001). Although the primer efficiency for *Ago2* were 127% (Table S4), amplification specificity was validated with melt-curve analysis and no-template controls. We therefore interpreted knockdowns based on the direction and statistical significance of expression changes rather than absolute fold-change values.

Small RNA libraries were prepared from a pool of three Ago2-dsRNA treated mites that exhibited the highest relative knockdown, and three randomly selected GFP-dsRNA control mites, with the methods described above. These samples were uploaded to the Genbank Sequence Read Archive under the BioProject PRJNA1470040 (Accessions: SRR38816849 and SRR38816850).

## 3 RESULTS

### 3.1 Small RNA composition in *V. destructor* mites

*Varroa destructor* small RNA libraries were comprised of 26.3%–44.26% of reads mapping to the *V. destructor* genome, 0.82%–1.63% of reads mapping to the *A. mellifera* genome, and 12.68%–27.05% of reads mapped to both genomes. Between the individual libraries, the percentage of reads mapping to the curated New Zealand virus reference library (Table S2) varied from 14.9%–42.16%, and 10.43%–19.48% were left unannotated (Figure 1A). We identified no novel viruses with BLASTn and DIAMOND homology searches against NCBI’s nucleotide and protein databases, respectively.

**Figure 1.**
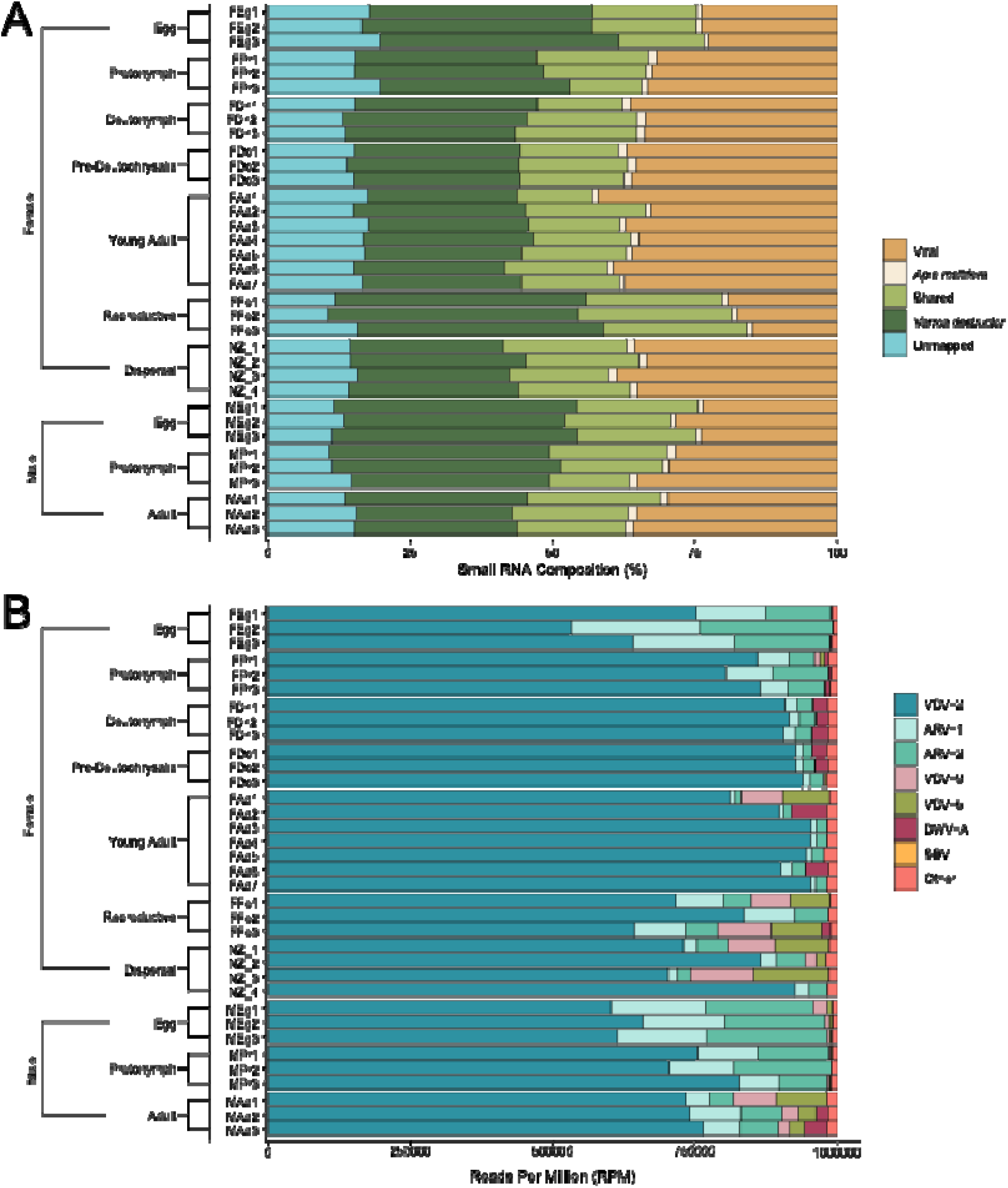
Small RNA read composition from *Varroa destructor* life-stages. Each row represents individual mite samples (n = 35), life stage and sex is indicated on the left. **(A)** Small RNA reads were mapped to the *V. destructor* genome (Vdes_3.0, Genbank Accession: GCA_002443255.1), then to the *A. mellifera* genome (Amel_HAv3.1, Genbank Accession: GCA_003254395.2), and finally to the viral reference library (Table S2). “Shared” represents reads that mapped to both genomes. The “Unmapped” portion represent reads that did not map to any of the aforementioned genomes and libraries. **(B)** Viral composition derived from reads mapping to the “Viral” sRNA portion of each sample (RPM-normalised). Viruses were included if detected in at least one sample, with > 100 reads, >30% genome coverage, and an RPM >1. Viruses that did not meet these thresholds were grouped with other unknown reads as “Other”.

The reads mapping to the viral reference library were defined as virus-derived siRNA (vsiRNA). A virus was retained in the dataset if at least one sample contained greater than 100 mapped reads, covered over 30% of the genome, and if the RPM exceeded 1. There were multiple DWV strains in the viral reference library (DWV-B, DWV-C, DWV-D, and Kakugo virus) which all share greater than 79% homology to the DWV-A_NZ genome, resulting in similarity across the short (<50 nt) vsiRNA mapped reads, leading to spurious alignments with sufficient read counts and genome coverage to pass the inclusion criteria. Visual inspection confirmed the presence of other DWV strains to be unlikely due to short, interspersed coverage over regions with homology to the DWV-A_NZ strain, so reads mapping to all other DWV strains were therefore combined with the DWV-A_NZ read counts. A total of seven viruses passed filtering criteria, including two -ssRNA viruses, Apis Rhabdovirus 1 (ARV-1), and Apis Rhabdovirus 2 (ARV-2); and five +ssRNA viruses, Varroa destructor virus 2 (VDV-2), Varroa destructor virus 5 (VDV-5), Varroa destructor virus 9 (VDV-9), Deformed wing virus A (DWV-A), and Sacbrood virus (SBV) (Figure 1B). ARV-1, ARV-2, and VDV-2 were the most prevalent viruses, detected in all *V. destructor* life stage samples, followed by DWV-A which was detected in all samples except for two female young adults (W12 and W16). VDV-2 was the most abundant virus, ranging from 531,683–954,547 RPM, followed by ARV-1 and ARV-2 which represented 7,605–225,782 RPM and 11,155–234,108 RPM, respectively.

### 3.2 Identification of eight co-infecting variants of Varroa destructor virus 2

To better capture the complete proportion of vsiRNA reads belonging to the highly diverse VDV-2, we assembled *de novo* eight distinct VDV-2 strains from New Zealand *V. destructor* transcriptomes (Lester, Felden et al. 2022), with each strain denoted as -alpha, -beta, -gamma, -delta, -epsilon, -theta, -zeta and -eta. Each assembled VDV-2 strain encompassed between 5,200–9,100 nt of the VDV-2 genome and exhibits 74-90% pairwise identity with the VDV-2 reference strain (Israel; NC_040601; Table S2). We observed a similar level of diversity in VDV-2 sequences assembled from *V. destructor*life stage samples from the US, with each of the eight US variants showing pairwise identities of 93-98% with their corresponding NZ variant. All eight variants from both locations have <91% nucleotide identity to the three previously published VDV-2 genome sequences.

VDV-2 strains from NZ and the US showed strong phylogenetic clustering as demonstrated by maximum likelihood analysis, with six of the eight variants clustering as distinct pairs (-alpha, -beta, -gamma, -delta, -epsilon, -theta); while two (-zeta and -eta) formed a separate mixed clade (Figure 2A). When compared to global reference sequences, the eight strains separate into two clear clades: a VDV-2-China-like clade comprised of the -epsilon, -zeta, -eta and -theta strains; and a VDV-2-UK-like clade containing -alpha, -beta, -gamma and -delta strains alongside the VDV-2 reference from Israel and two *Varroa jacobsoni* VJV-2 isolates from PNG.

**Figure 2.**
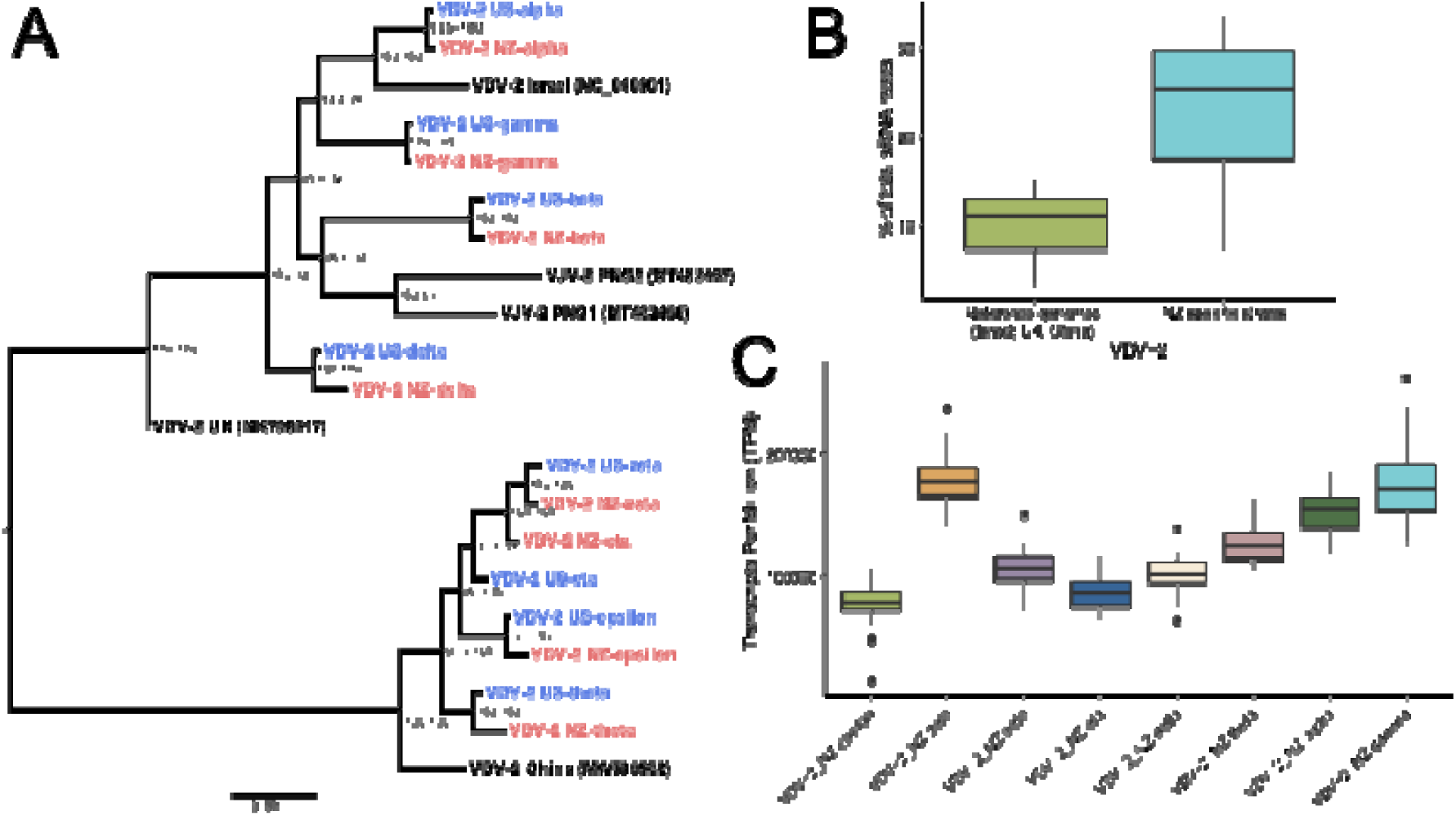
Varroa destructor virus 2 (VDV-2) strain diversity. **(A)** Maximum likelihood phylogenetic tree of each of the eight VDV-2 strains assembled from New Zealand (red) and US (blue) *V. destructor* transcriptomes, compared to five available VDV-2 sequences obtained from NCBI (black). Nucleotide sequences were aligned using MUSCLE and trimmed for gaps and incomplete sequences, leaving a final length of 3950 nucleotides. The phylogenetic tree was generated using maximum likelihood in IQ-TREE with the best-fit substitution model of TIM2+F+I+G4, which had the optimal BIC score as determined by ModelFinder. Branch supports were estimated using Ultrafast bootstrap approximation [UFBoot (Hoang et al., 2018)] using 1,000 replicates. Support values shown are SH-aLRT and UFBoot supports. **(B)** Proportion of total sRNA reads mapping to available VDV-2 sequences from NCBI (green), compared to the eight NZ-specific VDV-2 strains (blue) in all *V. destructor* life stage samples (n = 35). **(C)** Viral abundance of the eight VDV-2_NZ-specific strains across all *V. destructor* life stage samples. Abundance expressed as transcript per million (TPM) of viral siRNA reads, accounting for viral genome length. Strains ordered by assembled contig size from smallest (left) to largest (right). All eight VDV-2 strains are present in all samples

The use of NZ-specific strains significantly improved the proportion of vsiRNA reads mapping to VDV-2 in all mite samples (paired Wilcoxon signed-rank test, *V* = 630, *p* < 0.001, n = 35), with the mean percentage of total sRNA reads mapping to VDV-2 increasing from 10.1% (reference genomes) to 23.3% (Figure 2B), indicating that single mites host a diverse mix of VDV-2 strains. When normalised by contig length, VDV-2-zeta was the most abundant, followed by gamma, alpha, theta, beta, delta, eta, and epsilon (Figure 2C).

### 3.3 *V. destructor* life stage drives viral communities

Six viruses were prevalent among all *Varroa*’s life stages: DWV-A, VDV-2 (all eight strains), ARV-1, ARV-2, VDV-5, and VDV-9, whereas SBV was only found in eggs and protonymphs (Figure 3A). VDV-2 was consistently the most abundant virus amongst all life stages. ARV-1 and ARV-2 abundance was highest in eggs and protonymphs with reduced levels in other life stages. VDV-9 and VDV-5 abundances were higher in adults and foundresses and exhibited lower levels in immature *V. destructor* life stages. Interestingly, DWV-A_NZ was low in eggs and mature adults, exhibiting higher abundance in deutonymphs, pre-deutochrysalises, and young adults. Hierarchical clustering of the vsiRNA abundances across the *V. destructor* life stages separated younger life stages from older life stages (Figure 3A). Viruses clustered based on shared abundance patterns, with ARV-1 clustering with ARV-2, and VDV-9 and VDV-5 clustering together.

**Figure 3.**
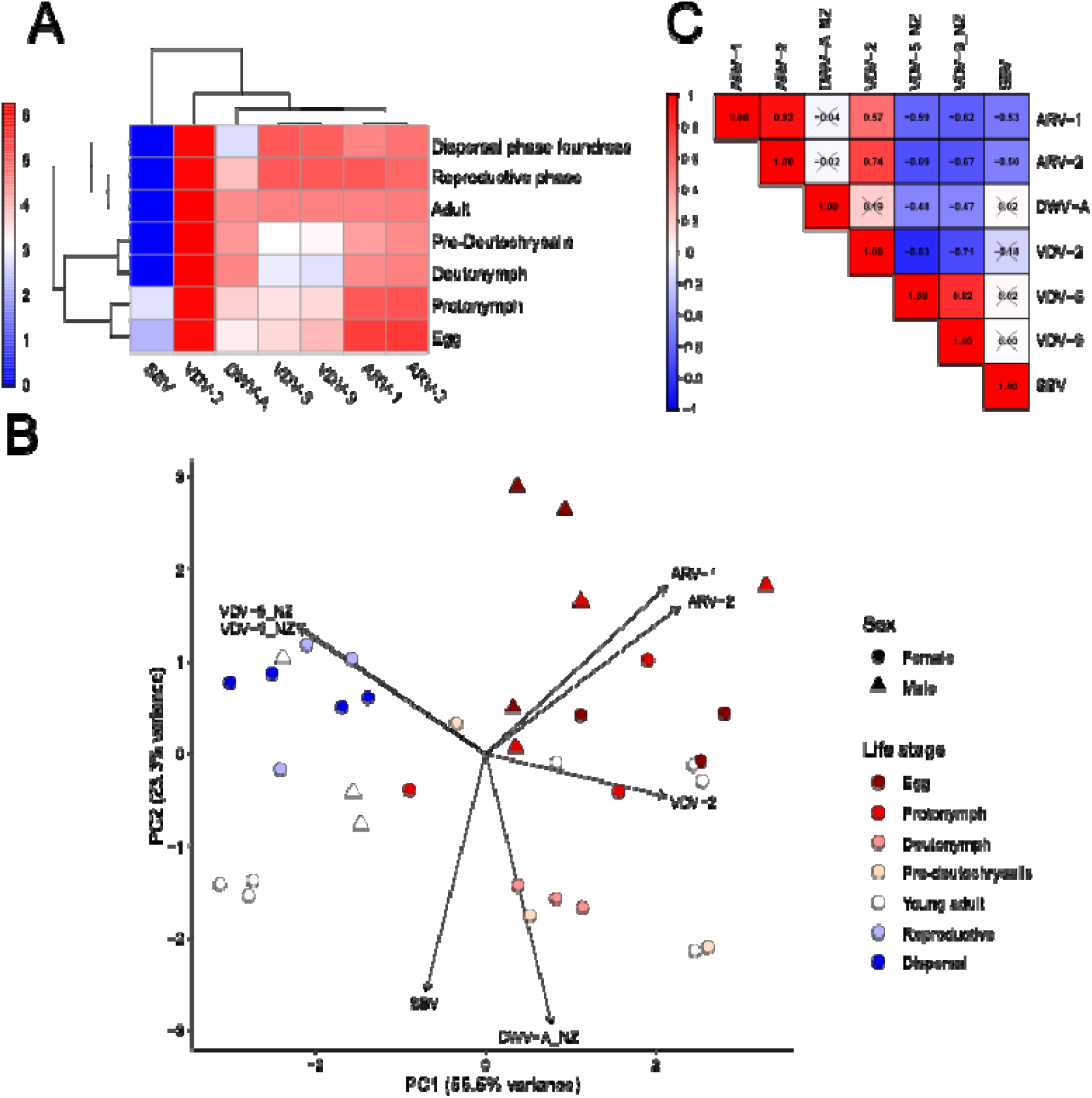
*Varroa destructor* life stage viral communities and correlation networks. **(A)** Abundances of viruses from viral read counts across all replicates. Counts were normalised to reads per million (RPM), and log10-transformed. Rows and columns are hierarchically clustered using Euclidean distances. **(B)** Pairwise correlations among viruses calculated using Spearman’s rank correlation. Correlation shown in each cell with the colour indicating the magnitude. Non-significant correlations are marked with a cross. **(C)** Principal component analysis of viral abundances derived from CLR-transformed viral RPM values, using Euclidean distances. Life stages are colour coded and sex are denoted by shape (females = circles; males = triangles). Arrows represent PCA loading vectors, indicating the direction of each virus’s contribution to the two principal components. Principal components 1 and 2 account for 78.8% of the variance.

Similar to the hierarchical clustering, PCA ordination of CLR-transformed, RPM normalised vsiRNA reads revealed distinct clustering among the life stages. The first two principal components explained 78.8% of variance (PC1 = 55.5%, PC2 = 23.3%; Figure 3B). Along PC1, younger life stages (eggs to pre-deutochrysalis) clustered separately from the older life stages (adult and foundresses), with exception of young female adults which exhibited overlap across these two clusters. Loading vectors indicated that ARV-1 and ARV-2 were associated with early life stages, particularly eggs and protonymphs; VDV-5 and VDV-9 are associated older life stages in adults and foundresses; and DWV-A showing the strongest association to intermediate life stages in deutonymphs. Viruses clustered along PC2 according to host range, distinguishing bee-associated viruses (DWV-A and SBV) from *Varroa*-associated viruses (ARV-1, ARV-2, VDV-2, VDV-5, and VDV-9).

To further assess the viral community differences across the different life stages and between sexes, a two-way PERMANOVA of Bray-Curtis dissimilarities generated from RPM-normalised vsiRNA reads was conducted. Life stage and sex together explained 75.7% of the variation in viral community composition, with life stage accounting for the majority of the variation (*R^2^* = 0.68, *F*_6,27_ = 12.56, *p* < 0.001), whilst sex contributing to a smaller but significant component (*R^2^* = 0.08, *F*_1,27_ = 8.78, *p* < 0.002).

### 3.4 Distinct viral correlation networks across *V. destructor* life stages

The Spearman’s correlation plot of the CLR-transformed vsiRNA RPM values across *V. destructor* life stages revealed clear patterns of viral prevalence (Figure 3C). ARV-1 and ARV-2 were significantly correlated with each other (*R* = 0.93, *p* < 0.001). VDV-2 exhibited a stronger positive correlation to ARV-2 (*R* = 0.74) than with ARV-1 (*R* = 0.57); and DWV-A was not significantly correlated with ARV-1, ARV-2, and VDV-2 (Figure 3C). VDV-5 and VDV-9 abundances were significantly positively correlated (*R* = 0.82, *p* < 0.001), but negatively correlated with ARV-1, ARV-2, VDV-2, and DWV-A. SBV showed only a significantly negative correlation with ARV-1 and ARV-2 (*R* = -0.53 and *R* =-0.50, *p* < 0.001 respectively), whilst exhibiting non-significant abundance relationships with all other viruses (Figure 3C).

### 3.5 vsiRNA profiles reveal *V. destructor*’s core virome

Five viruses exhibited vsiRNA patterns that are consistent with RNAi-mediated degradation signals across all *V. destructor* life stages and all individual samples, with greater than ∼90% of reads at antisense polarity and the majority of reads distributed at 24 nt (Figure 4). These viruses include two -ssRNA viruses (ARV-1 and ARV-2; Figure 4A, B) and three +ssRNA viruses (VDV-2, VDV-5, and VDV-9; Figure 4C-E). Additionally, RNAi-mediated vsiRNA degradation profiles were evident for all eight VDV-2 strains in all samples (Figure S1). As expected, a lower proportion of sense reads was observed for the two -ssRNA viruses present (ARV-1 and ARV-2), typically comprising less than 1% of vsiRNA reads, in contrast to +ssRNA viruses, which ranged from 3.21-9.70% of vsiRNA reads, with a 23 nt sense peak. In the three samples positive for SBV, sRNA reads mapping to the SBV genome displayed signals of non-specific degradation, with no clear vsiRNA profiles (Figure S2).

**Figure 4.**
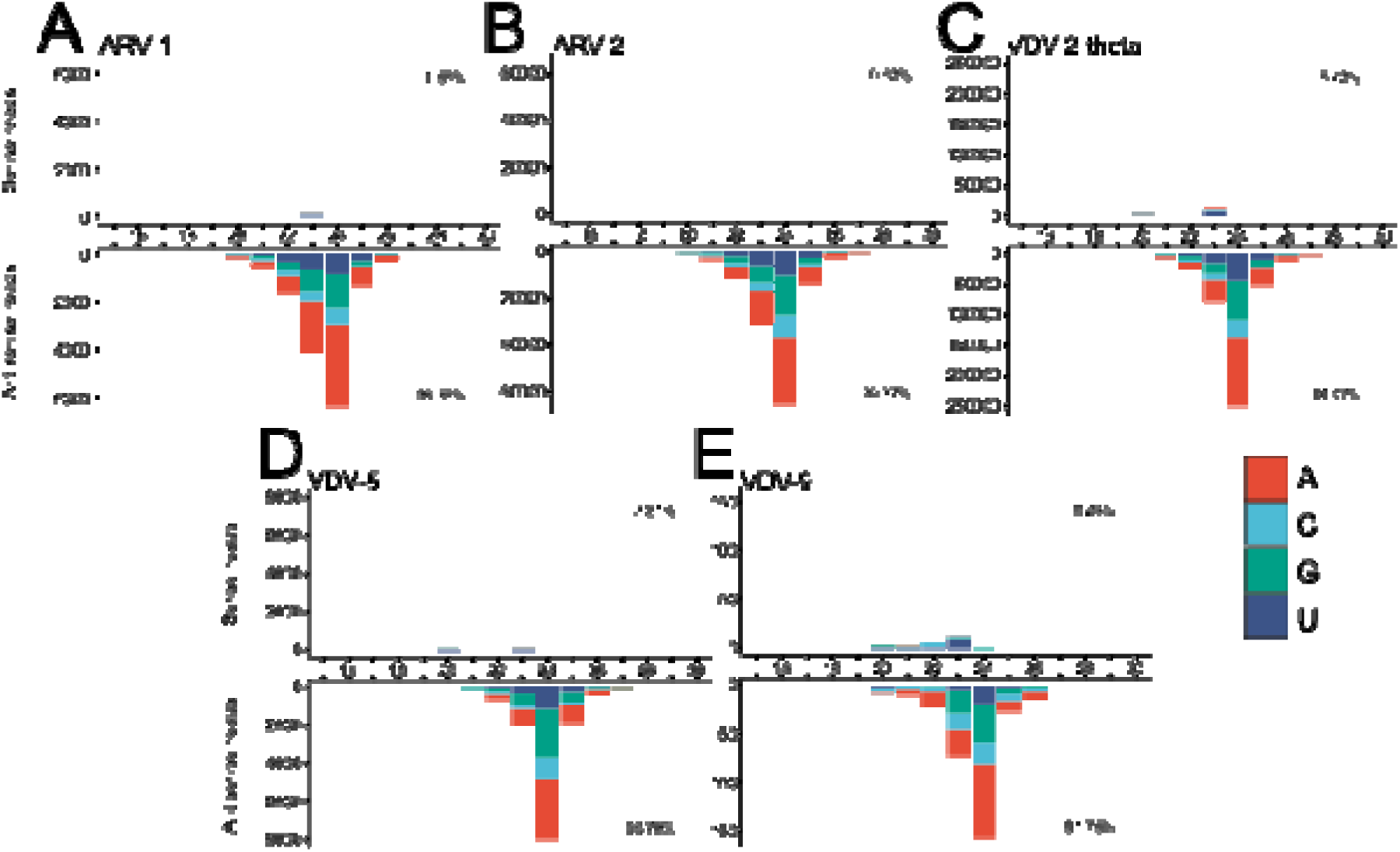
Virus-derived siRNA (vsiRNA) profiles observed in *V. destructor* life stages. vsiRNA profiles of reads mapping to **(A)** Apis rhabdovirus 1 (MF114351); **(B)** Apis rhabdovirus 2 (MZ821796); **(C)** Varroa destructor virus 2 (theta strain; PZ380883); **(D)** Varroa destructor virus 5 (PZ380885); **(E)** Varroa destructor virus 9 (OR224325). Representative profiles were selected at random (A = FAd2; B = FDc2; C = FPr2; E = MEg2; E = FEg2). Read lengths (15-30 nt) depicted on the *x-*axis, and the first nucleotide of each mapped vsiRNA read is colour coded. A similar profile was observed at every life stage for each virus. The complete set of vsiRNA profiles for all samples and viruses are displayed in Figure S1, 4-7. F

### 3.6 Variation in DWV-A RNAi-mediated degradation across *V. destructor* life stages

Unlike other *Varroa*-infecting viruses, mites infected with DWV-A produced variable vsiRNA profiles across life stages and individuals. In DWV-A infected eggs and male protonymphs (samples FEg2, MEg1, MEg2, MEg3, and MPr2), vsiRNA profiles had clear sense and antisense 23 nt peaks, along with a range of non-specific fragments with sense polarity (Figure 5A, S3). In contrast, most intermediate life stages (3 pre-deutochrysalises, 2 deutonymphs, and 2 young adults) produced RNAi-mediated degradation signals with a high proportion of antisense 24 nt fragments, consistent with the characteristic *Varroa* antiviral response to a replicating virus (Figure 5B, S3). Finally, in older DWV-A infected mites (reproductive foundresses and dispersal females) we also observed antisense 24 nt peaks but with a higher proportion of non-specific, sense degradation fragments (Figure 5C, S3).

**Figure 5.**
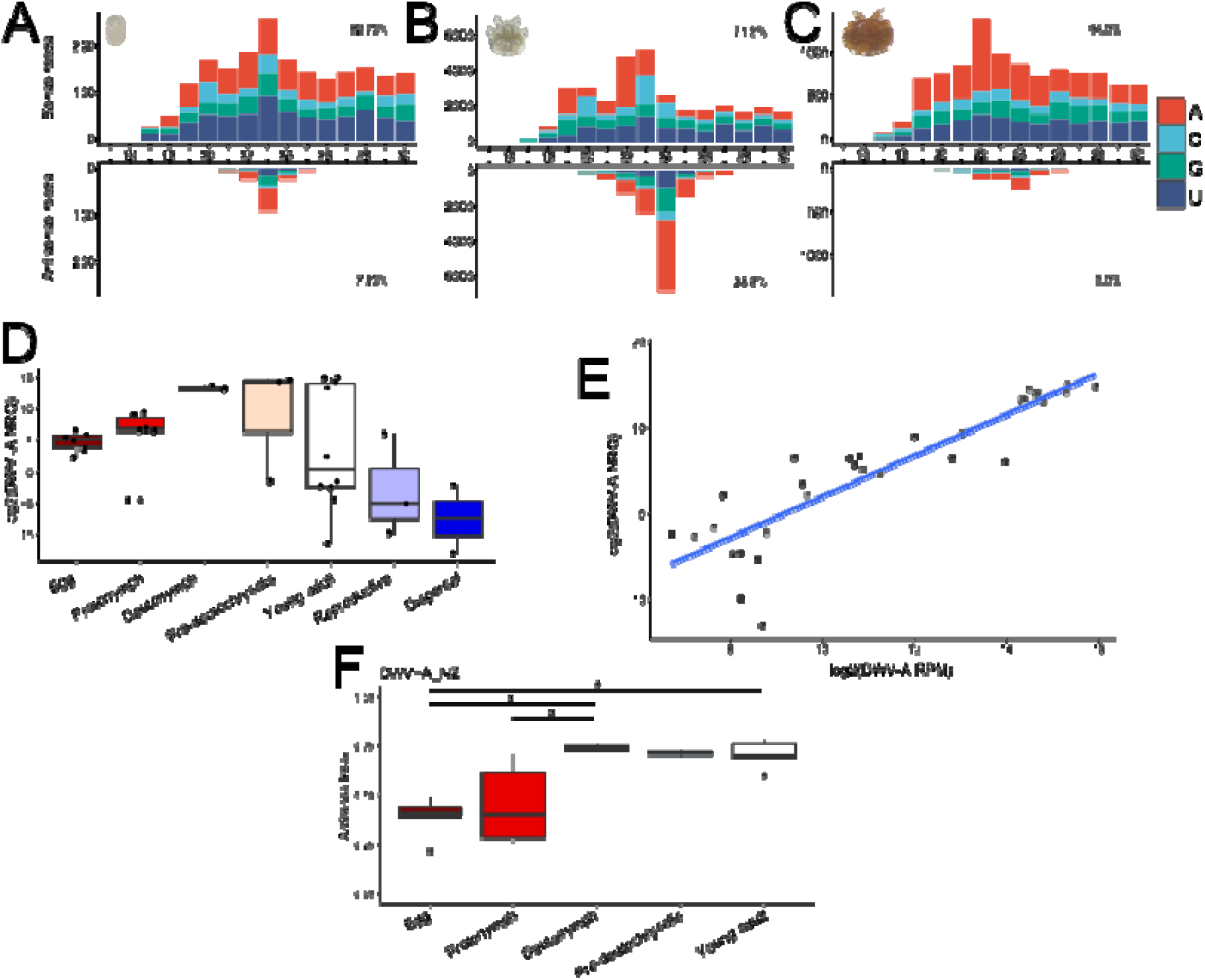
vsiRNA profiles of small RNA reads mapping to the DWV-A genome (MN538208.1) in **(A)** immature (egg; FEg2), **(B)** intermediate (pre-deutochrysalis; FDn2), and **(C)** adult life stages (Female reproductive foundress; FFo3. All DWV-A vsiRNA profiles are displayed in Figure S3. **(D)** Quantitative PCR comparing DWV-A viral load across mite life stages calculated from log2transformed normalised relative quantities (NRQ), using the geometric mean of two reference genes (18S and NADHD). **(E)** Correlation between log2 DWV-A RPM from small RNA and log2 DWV-A NRQ from qPCR. Blue line indicates linear regression (Pearson’s r = 0.89, p < 0.001) and shaded area shows 95% CI. **(F)** DWV-A vsiRNA antisense index reflecting the dominant antisense fragment size within each vsiRNA profile (between 23-24nt). Higher values represent greater proportion of 24 nt antisense reads. Statistical significance assessed with Kruskal-Wallis and Dunn’s post-hoc tests, with adjusted p-values using the false discovery rate (FDR). Significant differences represented with an asterisk. Antisense indices were lowest in eggs and protonymphs.

To determine whether these differences in vsiRNA levels reflect variation in DWV-A genome abundance throughout *V. destructor* life stages, we examined DWV loads using qPCR. Life stage accounted for a substantial proportion of the variance (η^2^ = 0.36), with viral loads showing a trend towards higher loads in intermediate life stages (ANOVA *F*_6, 26_ = 2.46, *p* = 0.051; Figure 5D). DWV vsiRNA abundance was strongly positively correlated with qPCR-derived relative DWV-A viral loads in all samples (Pearson’s *r* = 0.89, *df* = 29, *p* < 0.001; Figure 5E), indicating that vsiRNA abundance broadly reflects the relative DWV-A genome levels.

The antisense index, comparing the proportion of 23 nt sense vs 24 nt antisense vsiRNAs, differed significantly among life stages for DWV-A-infected mites (Kruskal-Wallis χ*^2^* = 13.046, df = 4, *p* = 0.011; Figure 5F). Early developmental stages (eggs and protonymphs) exhibited lower antisense indices than later life stages (deutonymphs, pre-deutochrysalis and adults), indicating a higher proportion of 23 nt fragments, with post-hoc Dunn’s tests (FDR-corrected) revealing significant differences between eggs and both deutonymphs and adults, and between protonymphs and deutonymphs (all *p*-adjusted < 0.05). The DWV-A loads in dispersal and reproductive foundress life stages were low, so that the vsiRNAs did not meet the minimum antisense read count threshold for calculation and were excluded from the analysis. The antisense index in pre-deutochrysalis indicated similarly elevated 24 nt vsiRNAs, however, the small number of DWV-infected samples (n = 2) likely limited statistical power to detect pairwise differences following multiple-testing correction.

### 3.7 Evidence that primary and secondary siRNAs are produced by processing of exogenous dsRNA

Since invertebrate RNAi is triggered by dsRNA produced by replicating viruses, we used synthetically produced dsRNA to investigate the processing steps involved in this antiviral response. We treated *Varroa* with 800bp dsRNA targeting GFP, to measure the primary processing of exogenous dsRNA, and 800bp dsRNA targeting one of *V. destructor*’s *Ago2* homologues (*LOC111255253*) to identify secondary siRNA responses to an expressed gene.

In GFP-dsRNA treated mites, 12.01% of all small RNA reads mapped to the 800 bp dsRNA construct, with an average read depth of 2188.14 (Figure 6A). The GFP siRNA profiles showed clear 23 nt sense and antisense fragment peaks (Figure 6B) of roughly equal abundance, and a low antisense ratio of 0.25, similar to the antisense ratios observed for DWV-A in eggs, suggesting Dicer-mediated primary siRNA processing of the GFP-dsRNA, likely by a Dicer2 analogue.

**Figure 6.**
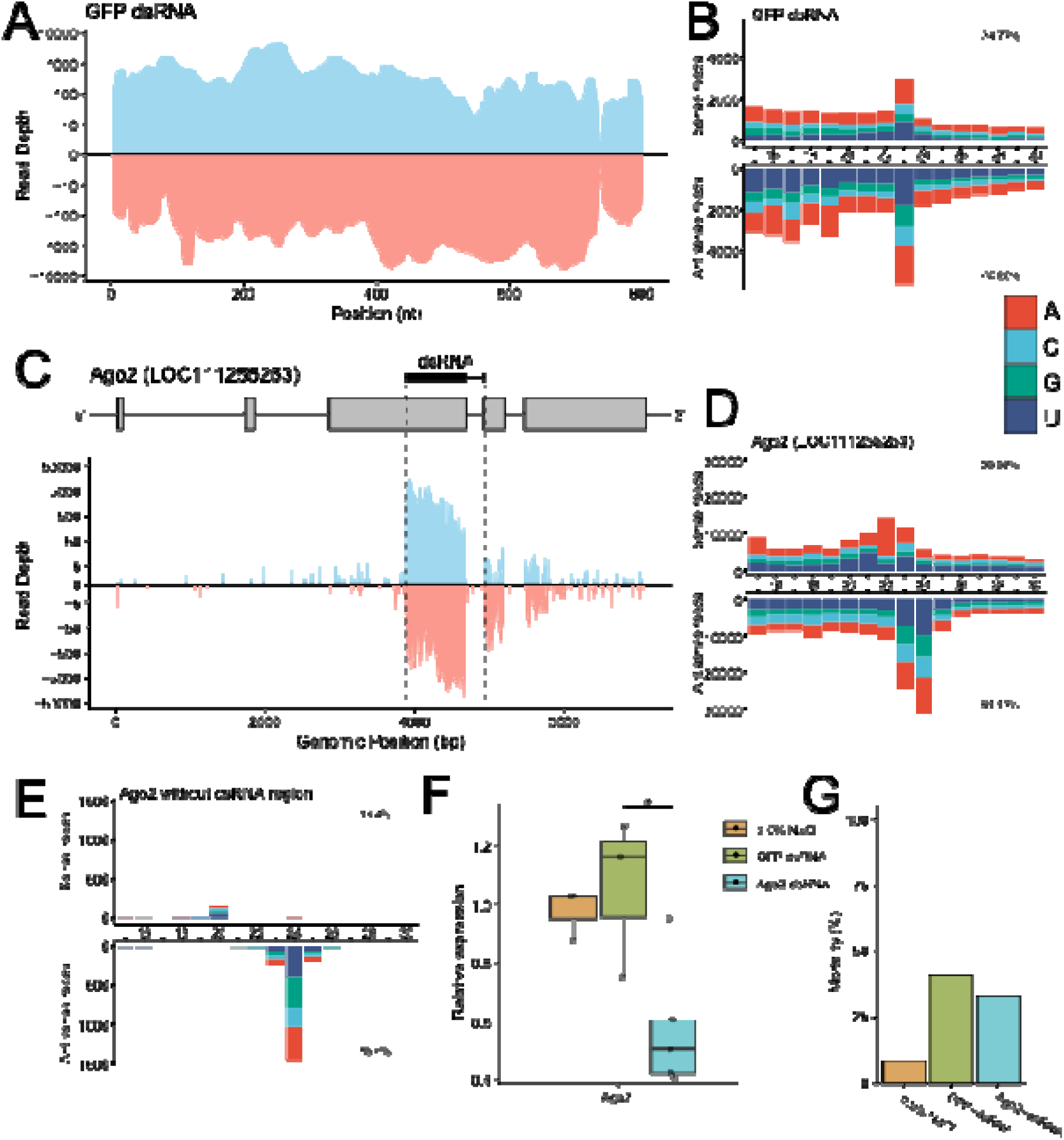
Exogenous dsRNA processing in *V. destructor*. Small RNA read depths of **(A)** the GFP sequence, and **(C)** the *Ago2* locus (*LOC111255253*), following dsRNA treatment with GFP-dsRNA and Ago2-dsRNA, respectively. The *x*-axis depicts the position along sequence, while the *y*-axis shows the read depth (pseudo-log_10_ transformed). Sense reads are shown above the *x*-axis in blue, while antisense reads are below in red. For the *Ago2* locus, exons are indicated by grey boxes, and the dsRNA region by the dotted lines. Notably, the sRNA reads extend downstream of the dsRNA target region. Small interfering RNA (siRNA) profiles of reads mapping to the **(B)** GFP sequence, **(D)** *Ago2* locus, and **(E)** the *Ago2* locus without the dsRNA target site. **(F)** Relative expression of *Ago2* in non-specific 557 dsRNA (GFP) or Ago2-dsRNA treated mites, compared to 0.9% NaCl controls. Relative expression was calculated with the Pfaffl method, normalised to two endogenous reference genes NADHD and 18S (Table S4). Data were analysed using a one-way ANOVA (F_2, 8_ = 5.92, p < 0.05), followed by pairwise Tukey’s HSD post-hoc tests. Significant differences are represented with an asterisk. **(G)** Mortality of mites treated with exogenous dsRNA. Mites were exposed to 0.9% NaCl, GFP563 dsRNA and Ago2-dsRNA overnight, then allowed to recover on pupae for 48 hours.

Mapping of the sRNA reads to the *Ago2* locus revealed a large accumulation of reads within the 800 bp dsRNA recognition site, with an average read depth of 9282.2 compared to 9.14 in the flanking regions (Figure 6C). To ensure the specificity of the dsRNA construct, the 800 bp Ago2-dsRNA target region was designed to encompass an exon-exon junction. Only 12 small RNA reads mapped to the corresponding intronic region, indicating dsRNA construct specificity for the mature *Ago2* mRNA transcript (Figure 6C). In contrast to the GFP-dsRNA siRNA profiles, mites treated with Ago2-dsRNA produced siRNA profiles dominated by both 24 and 23 nt fragments, representing 11.69% and 11.50% of all reads, respectively (Figure 6D). Analysis of reads mapping to the flanking regions outside of the Ago2-dsRNA site revealed a distinct siRNA profile with clear 24 nt antisense bias (Figure 6E), identical to the profiles we observed for actively replicating viruses, suggesting a secondary siRNA response leading to RNAi degradation of the full *Ago2* mRNA transcript. Furthermore, this secondary response showed a distinct 3’ bias (Figure 6C), with read depth downstream the Ago2-dsRNA target site (24.4-fold) substantially higher than that upstream of the region (1.0-fold), suggesting the secondary siRNA response spreads downstream from the primary trigger site.

To confirm that the activation of the RNAi pathway resulted in gene knockdown, we measured transcript abundance by qRT-PCR and found that Ago2-dsRNA treatment resulted in significantly decreased *Ago2* expression compared to GFP controls (Figure 6F; ANOVA F_2, 8_ = 5.92, p < 0.05; Tukey’s HSD, p = 0.035), with a similar but less pronounced trend observed when compared to NaCl controls (p = 0.078). Treatment with GFP-dsRNA and Ago2-dsRNA resulted in mortality rates of 42% and 33.3%, respectively, compared to 8.3% mortality of control treatments (Figure 6G).

## 4 DISCUSSION

This work helps to establish vsiRNA profiling as a powerful tool for dissecting virus transmission and co□infection dynamics in arthropods. By analysing the unique vsiRNA profiles produced by *V. destructor*, we characterised viral transmission and co-infection dynamics throughout mite development. Based on vsiRNA profiles, we identified six actively replicating viruses in our population of *V. destructor*. The antiviral response to five of these viruses (ARV-1, ARV-2, VDV-2, VDV-5 and VDV-9) shows a typical *Varroa* profile of 24 nt antisense vsiRNAs during all life stages, suggesting viral infections are already established in embryos. However, the response to the sixth actively replicating virus, DWV-A, shows a different profile in early life stages, with sense and antisense 23 nt vsiRNAs in eggs and protonymphs, transitioning to the 24 nt antisense vsiRNA profile in deutonymphs and later developmental stages. The change in vsiRNA fragment size during development indicates that DWV is acquired and processed differently to the other five viruses in younger mites, while still clearly showing signatures of replication. We propose that the vsiRNA size-switch between early infection (23 nt sense/antisense) to established infection (24 nt antisense) indicates the presence of primary and secondary siRNA responses in *V. destructor.* The primary siRNA response consists of Dicer-produced, sense-antisense vsiRNAs when viral dsRNA is first encountered, followed by a shift to a secondary, antisense-biased vsiRNA profile as typical for *V. destructor* as the virus infection becomes established.

While our previous studies of vsiRNA in *V. destructor* have demonstrated a strong bias for 24 nt antisense vsiRNA (Damayo et al., 2023; Remnant et al., 2017), our observation of 23 nt sense-antisense DWV-A vsiRNA in early life stages provides evidence that in *Varroa*, the antiviral-RNAi immune pathway involves multiple hierarchical processing steps. Sense-antisense siRNA duplexes are typically generated from the enzymatic cleavage of long dsRNA by Dicer-2 (Zamore et al., 2000). Across eukaryotes, siRNAs range in size from 21-28 nt (MacRae et al., 2007; Siomi & Siomi, 2009). Insects typically produce 21-22 nt duplexes (Elbashir et al., 2001), whereas arachnids possess siRNAs that are more variable in size. Tick species such as the black-legged tick (*Ixodes scapularis)* and Asian long-horned tick (*Haemaphysalis longicornis)* generate 22 nt sense-antisense vsiRNA profiles *in vitro* (Schnettler et al., 2014) and *in vivo* (Xu et al., 2021), in contrast to the larger, 23 nt DWV-A-derived vsiRNA duplexes we observed during early infection in *Varroa*. The 23 nt vsiRNA duplexes are likely to be produced by one or more of *Varroa*’s multiple *Dicer-2* homologues (Nganso et al., 2020), consistent with the primary biogenesis of siRNA duplexes in other organisms. When a virus begins to replicate at sufficient levels and an infection becomes established, a 24 nt antisense response emerges, suggesting secondary siRNA amplification potentially mediated by an *RdRP* homologue. The *V. destructor* genome contains an expanded repertoire of RNAi genes implicated in both primary and secondary siRNA biogenesis pathways, including at least two *Dicer2* and four *RdRP* homologues (Nganso et al., 2020), that may facilitate the production of these two differing vsiRNA profiles. The ancestral eukaryote likely had three ancestral *RdRP* paralogues, which were all lost in most animal taxa, with the exception of a few invertebrate lineages, such as lancelets, nematodes, and chelicerates (Lewis et al., 2018; Pinzón et al., 2019; Zong et al., 2009).

To distinguish primary dsRNA processing from secondary siRNA amplification, we exposed *V. destructor* to exogenous dsRNA targeting a non-expressed transcript, GFP. Much like the DWV-A vsiRNA profiles in younger *V. destructor* life stages, GFP-dsRNA was processed into 23 nt sense-antisense fragments. GFP-dsRNA treatment resulted in high mortality rates in *V. destructor*, that were consistent with observations in other dsRNA immersion studies (Becchimanzi et al., 2020), and oral acquisition via dsRNA-fed honey bees (McGruddy et al., 2024), raising the question of whether treatment with exogenous dsRNA is inherently toxic to *V. destructor*. Because GFP transcripts are not expressed in *Varroa*, we assume that secondary siRNA responses cannot occur due to the lack of an mRNA template that RdRPs require for amplification (Pak & Fire, 2007; Wassenegger & Krczal, 2006), in line with the lack of 24 nt antisense GFP fragments observed. Conversely, exposure to a dsRNA construct targeting an expressed gene (*Ago2)* revealed evidence of secondary siRNA amplification. We observed enrichment for both 24 nt and 23 nt siRNA within the Ago2-dsRNA target site, and notably, an accumulation of 24 nt antisense siRNAs downstream of the dsRNA construct in the 3’ flanking region. Antisense biased siRNA are hallmarks of RdRP synthesis in nematodes, however in *C. elegans*, the mechanism of secondary siRNA synthesis occurs via 5’ transitivity (the processive production of secondary siRNAs in a 3’→5’ direction), yielding 22 nt antisense small RNA possessing 5’ triphosphates (Pak & Fire, 2007; Sijen et al., 2001). In *V. destructor,* the prevalence of 3’ secondary siRNA suggests a mechanism that diverges from 5’ transitivity observed in *C. elegans*, which may be conserved amongst Chelicerata; for instance, *I. scapulari*s ISE6 cells have been shown to produce RdRP-dependent small RNAs that are similarly enriched in the 3’ UTR of target genes (Feng et al., 2023). Functional characterisation of *V. destructor’s* complete RNAi repertoire is required to determine whether secondary siRNA synthesis relies on RdRP priming with primary siRNA like *C. elegans* (Aoki et al., 2007), or is synthesised *de novo* from sliced target transcripts akin to plant secondary siRNAs (Yoshikawa et al., 2005).

In *Varroa*, established infections in eggs are indicated by a strong secondary siRNA response (24 nt, antisense vsiRNAs). The highly abundant 24 nt antisense vsiRNA profiles observed for ARV-1, ARV-2, and VDV-2 in all life stages suggests that these viruses make up a core virome and are transmitted vertically. In addition, VDV-5 and VDV-9 show evidence of replication throughout development, though with reduced vsiRNA abundance in intermediate (deutonymph and pre-deutochrysalis) life stages. The transmission pathway for these five core *Varroa*-infecting viruses is most consistent with transovarial transmission via germline infection, given the ubiquitous infection of all the *V. destructor* life stages including eggs (Amiri et al., 2018; Lange et al., 2024). Co-occurrence of ARV-1 and ARV-2 is consistent with previous reports of simultaneous detection and shared abundance patterns, suggesting potential interdependence between these two viruses (Kadlečková et al., 2022). Interestingly, there is also a significant positive correlation in the abundance of vsiRNAs for VDV-5 and VDV-9, which are negatively correlated with the other replicating viruses. No significant correlations (positive or negative) have been observed with VDV-5 and other *V. destructor* viruses in previous transcriptomic and qPCR studies (Eliash et al., 2022; Herrero et al., 2019). Coordinated abundance patterns of VDV-5 and VDV-9 across life stages in our examined population of mites may reflect synergy between these two viruses. So far only a single RNA segment of the VDV-9 genome has been characterised from a predicted bipartite genome, which is distantly related to the capsid segment of other Beihai horseshoe crab virus 1 (Damayo et al., 2023). VDV-9 has exclusively been detected alongside either VDV-5 or VDV-3 (Damayo et al., 2023; Kim et al., 2026); two related viruses that share approximately 75% nucleotide identity (Herrero et al., 2019). Our consistent observations of mite co-infection with VDV-9 and VDV-5 could reflect a satellite-helper virus interaction, whereby the VDV-9 capsid segment is a subviral particle ‘satellite’ that depends on the replication machinery encoded by the ‘helper’ virus VDV-5 or, by extension, the closely related VDV-3 (Gnanasekaran & Chakraborty, 2018). Viral satellites can affect pathogenicity of their cognate helper virus; for example satellites can attenuate Euphorbia yellow mosaic virus accumulation and disease severity possibly by competing for encapsidation (Mar et al., 2017). However, as the symptoms of *Varroa*-infecting viruses are yet unknown, it is still unclear how the VDV-9/VDV-5 interaction, or the interaction between ARV-1 and ARV-2, could impact viral pathogenicity and *Varroa* biology more generally.

In addition to informing how viral processing, transmission and co-occurrence occurs in *V. destructor*, our small RNA sequence data has confirmed that the widespread Varroa destructor virus 2 is a highly diverse, multi-strain virus. Previously we observed high strain diversity in VDV-2 assembled *de novo* from *V. destructor* transcriptomes in NZ (Lester et al, 2022). Here we clarify the sequences of each strain and show that vsiRNAs map to each of the eight variants in significant loads in all mite life stages. The co-existence of eight strains in all individuals suggest that VDV-2 is stably vertically co-transmitted as a dynamic population of genetically distinct variants. The significance of VDV-2 multi-strain dynamics (Makau et al., 2022) on *Varroa* biology is unclear. Western honey bees (*Apis mellifera*) are similarly infected with the highly variable multi-strain Lake Sinai virus (Hou et al., 2023), which appears to be stably associated with the honey bee host, though unlike VDV-2 it is not common for individuals to be co-infected with such a high number of different strains. LSV strains are highly variable, with multiple diverse strains that coexist globally throughout honey bee populations (Daughenbaugh et al., 2015; Jones et al., 2026) rather than within individual hosts. The unique observation of VDV-2 diversity may provide an example of the evolutionary outcomes of the quasispecies theory (Lauring & Andino, 2010), where a single viral strain undergoes mutation and selection, leading to sufficient variation in each viral lineage such that the immune system encounters each differently. It is possible that *Varroa* has an established ‘germline lineage’ of viral strains that ensure ongoing viral survival and propagation (Lythgoe et al, 2017). It is also possible that individual cells are co-infected by multiple strains, leading to recombination and genomic reassortment, or that different strains occupy different cell types or exhibit tissue tropism, which could facilitate the high strain diversity observed. Further work is required to determine if VDV-2 variants share cellular co-infection or show tissue tropism.

Overall, our results provide evidence that DWV-A is one of many vertically transmitted viruses in *V. destructor*, though likely via a distinct mechanism from the germline transmission observed for the five other viruses. The 23 nt sense/antisense DWV-A vsiRNA profiles in eggs and nymphs are similar to GFP-dsRNA siRNA profiles, consistent with the Dicer-mediated processing of dsRNA substrates. The dsRNA from +ss RNA viruses arises from replication intermediates (Weber et al., 2006), and the presence of cellular dsRNA is necessary for Dicer-mediated production of the DWV 23 nt sense/antisense vsiRNA that we observe within eggs. It is unclear whether the DWV dsRNA present in eggs arises from viral replication, or passive carryover of DWV genomic RNA/dsRNA from foundress mites into eggs. DWV-A levels are low in foundress mites relative to eggs, suggesting that DWV-A RNA or virions are maternally acquired during oogenesis and are deposited into eggs, followed by replication and establishment of an active infection. It seems unlikely for *V. destructor* eggs to acquire DWV horizontally from developing pupae, given that foundresses lay eggs on the upper cell wall to avoid contact with the adjacent *A. mellifera* pupa, to minimise the risk of egg damage (Traynor et al., 2020).

In other arthropod systems, viral components can be incorporated into eggs by exploiting different oogenesis pathways. These include the hijacking of follicle cell vesicular trafficking pathways (Brasset et al., 2006), vitellogenin-virus complex formation for receptor-mediated endocytosis into oocytes (Huo et al., 2014), or by establishing persistent infections in the germ-line that persists through gametogenesis (Longdon & Jiggins, 2012). Remarkably, female *Varroa* (and other acarines) have a specialised somatic nutritive tissue called the lyrate organ (Sonenshine et al., 2022), implicated to be involved in kleptocytosis, a process where intact egg yolk precursor proteins and vitellogenins derived from *A. mellifera* parasitism are passed through the mite digestive system into the developing ova (Ramsey et al., 2022). It is possible that DWV-A particles or replicating RNA is transferred via kleptocytosis through the syncytial cytoplasmic continuity of the lyrate organ, either freely or bound by vitellogenins (Ramsey et al., 2022; Sonenshine et al., 2022). Given the prior detection of DWV proteins in *V. destructor* eggs (McAfee et al., 2017), together with our observation of primary vsiRNA responses in early life stages, it is plausible that either DWV-A viral particles or genomic RNA are transferred during oogenesis.

While vertical transmission of DWV-A from foundress to offspring may be feasible, it is unlikely to represent the primary transmission route for DWV-A in *V. destructor* populations. In honey bees, DWV-A can be transmitted vectorially from *V. destructor* (Posada-Florez et al., 2020), horizontally (de Miranda & Genersch, 2010; Posada-Florez et al., 2021; You et al., 2023), and vertically (Amiri et al., 2018). In parallel, our data suggest that DWV-A is transmitted vertically from *V. destructor* foundresses, and horizontally acquired through the ingestion of infected honey bee tissue, due to the accumulation of higher DWV-A loads in deutonymphs and pre-deutochrysalises – life stages characterised by higher host-feeding relative to body size when compared to adult mites (Han et al., 2024). The increase in DWV-A loads in deutonymphs and pre-deutochrysalises could occur due active viral replication, continual acquisition from honey bee hosts, or a combination of both.

By observing antiviral siRNA patterns across life stages, we demonstrate vertical transmission of the core *V. destructor* virome. In addition, we identified a potential satellite-helper virus interaction with VDV-9 and VDV-5, and clarified high strain diversity in VDV-2. Ultimately, our results suggest that *V. destructor* utilise primary and secondary siRNA pathways for antiviral defence, and our understanding of these pathways will impact the design, success and efficacy of emerging commercial RNAi-based biopesticides (McGruddy et al., 2024; Smeele et al., 2025). More broadly, this work highlights how vsiRNA profiling can provide powerful insights into virus transmission, persistence and co-infection dynamics in arthropods, where traditional approaches often struggle to resolve these processes.

## ACKNOWLEDGEMENTS

We thank Philip Lester’s Lab at Te Herenga Waka—Victoria University of Wellington for assistance with sample collection and laboratory support. This research was undertaken with the assistance of resources from the National Computational Infrastructure (NCI Australia), an NCRIS enabled capability supported by the Australian Government, and the technical assistance provided by the Sydney Informatics Hub, a Core Research Facility of the University of Sydney.

## CONTFLICT OF INTEREST

The authors declare no conflict of interest in relation to this work.

