## Supplementary materials for "Primary and secondary antiviral RNAi responses throughout *Varroa destructor* life stages reveal the vertical transmission of viruses"

**Table S1** *V. destructor* samples used in this study (n = 37). The number of individuals pooled in the sample is indicated in the column “n”.

| **Sample** | **Sex** | **Life stage** | **n** | **Study** | **SRA Accession** |
| --- | --- | --- | --- | --- | --- |
| MEg1 | Male | Egg | 4 | McKee et al. 2026 | SRR34672249 |
| MEg2 | Male | Egg | 4 | McKee et al. 2026 | SRR34672248 |
| MEg3 | Male | Egg | 4 | McKee et al. 2026 | SRR34672246 |
| MPr1 | Male | Protonymph | 4 | McKee et al. 2026 | SRR34672245 |
| MPr2 | Male | Protonymph | 4 | McKee et al. 2026 | SRR34672244 |
| MPr3 | Male | Protonymph | 4 | McKee et al. 2026 | SRR34672243 |
| MAd1 | Male | Adult | 4 | McKee et al. 2026 | SRR34672252 |
| MAd2 | Male | Adult | 4 | McKee et al. 2026 | SRR34672251 |
| MAd3 | Male | Adult | 4 | McKee et al. 2026 | SRR34672250 |
| FEg1 | Female | Egg | 2 | McKee et al. 2026 | SRR34672236 |
| FEg2 | Female | Egg | 4 | McKee et al. 2026 | SRR34672234 |
| FEg3 | Female | Egg | 4 | McKee et al. 2026 | SRR34672233 |
| FPr1 | Female | Protonymph | 3 | McKee et al. 2026 | SRR34672229 |
| FPr2 | Female | Protonymph | 4 | McKee et al. 2026 | SRR34672257 |
| FPr3 | Female | Protonymph | 4 | McKee et al. 2026 | SRR34672256 |
| FDn1 | Female | Deutonymph | 4 | McKee et al. 2026 | SRR34672255 |
| FDn2 | Female | Deutonymph | 4 | McKee et al. 2026 | SRR34672254 |
| FDn3 | Female | Deutonymph | 4 | McKee et al. 2026 | SRR34672253 |
| FDc1 | Female | Pre-deutochrysalis | 1 | McKee et al. 2026 | SRR34672259 |
| FDc2 | Female | Pre-deutochrysalis | 1 | McKee et al. 2026 | SRR34672258 |
| FDc3 | Female | Pre-deutochrysalis | 1 | McKee et al. 2026 | SRR34672247 |
| FAd1 | Female | Young adult | 1 | McKee et al. 2026 | SRR34672242 |
| FAd2 | Female | Young adult | 1 | McKee et al. 2026 | SRR34672241 |
| FAd3 | Female | Young adult | 1 | McKee et al. 2026 | SRR34672240 |
| FAd4 | Female | Young adult | 1 | McKee et al. 2026 | SRR34672239 |
| FAd5 | Female | Young adult | 1 | McKee et al. 2026 | SRR34672238 |
| FAd6 | Female | Young adult | 1 | McKee et al. 2026 | SRR34672237 |
| FAd7 | Female | Young adult | 1 | McKee et al. 2026 | SRR34672235 |
| FFo1 | Female | Reproductive phase foundress | 1 | McKee et al. 2026 | SRR34672232 |
| FFo2 | Female | Reproductive phase foundress | 1 | McKee et al. 2026 | SRR34672231 |
| FFo3 | Female | Reproductive phase foundress | 1 | McKee et al. 2026 | SRR34672230 |
| NZ1 | Female | Dispersal phase adult | 1 | Damayo et al. 2023 | SRR25010751 |
| NZ2 | Female | Dispersal phase adult | 1 | Damayo et al. 2023 | SRR25010750 |
| NZ3 | Female | Dispersal phase adult | 1 | Damayo et al. 2023 | SRR25010768 |
| NZ4 | Female | Dispersal phase adult | 1 | Damayo et al. 2023 | SRR25010767 |
| GFP-dsRNA | Female | Reproductive phase foundress | 3 | This study | SRR38816850 |
| Ago-dsRNA | Female | Reproductive phase foundress | 3 | This study | SRR38816849 |

**Table S2 –** Virus reference library containing viruses that have been reported in *V. destructor* and *A. mellifera*. Based on previously compiled viral reference libraries^1,2,3^, with New Zealand isolates used where available.

| Virus | accession |
| --- | --- |
| Anguilla anguilla circovirus | NC_023421.1 |
| Acute bee paralysis virus | NC_002548.1 |
| Aransas bay virus segment 1-6 | KC506162-KC506167 |
| Apis hypovirus 1 | MZ822101.1 |
| Apis hypovirus 2 | MZ822102.1 |
| Apis hypovirus 3 | MZ822103.1 |
| Apis iflavirus 1 | MZ822075.1 |
| Apis iflavirus 2 | MZ822076.1 |
| Apis mellifera filamentous virus | NC_027925.1 |
| Apis nora virus 2 | MZ822097.1 |
| Apis polycipivirus | MZ822098.1 |
| Apis picorna-like virus 1 | MZ822067.1 |
| Apis picorna-like virus 2 | MZ822068.1 |
| Apis picorna-like virus 3 | MZ822078.1 |
| Apis picorna-like virus 4 | MZ822079.1 |
| Apis picorna-like virus 5 | MZ822095.1 |
| Apis picorna-like virus 6 | MZ822096.1 |
| Apis rhabdovirus 1 | MF114351.1 |
| Apis rhabdovirus 2 | MZ821796.1 |
| Apis rhabdovirus 3 | MZ822104.1 |
| Apis rhabdovirus 4 | MZ822105.1 |
| Apis rhabdovirus 5 | MZ822106.1 |
| Antheraea pernyi iflavirus | KF751885.1 |
| Apis virga-like virus | MZ822100.1 |
| Bundaberg bee virus 1 | MG995701.1 |
| Bundaberg bee virus 2 | MG995700.1 |
| Bundaberg bee virus 3 | MG995702.1 |
| Bundaberg bee virus 4 | MG995705.1 |
| Bundaberg bee virus 5 | MG995706.1 |
| Bundaberg bee virus 6 | MG995707.1 |
| Bundaberg bee virus 7 | MG995703.1 |
| Bundaberg bee virus 8 | MG995704.1 |
| Bat circovirus | JF938080.1 |
| Bee Macula-like virus | KT162924.1 |
| Bat feces associated picorna-like virus, partial cds | JN857331.1 |
| Beihai horseshoe crab virus 1 | KX883115.1 |
| Black queen cell virus | NC_003784.1 |
| Chronic bee paralysis virus | NC_010711.1 |
| Cyclovirus NG14 | GQ404855.1 |
| Cyclovirus NGchicken8 | HQ738643.1 |
| Cyclovirus TN18 | GQ404858.1 |
| Cyclovirus TN25 | GQ404857.1 |
| Dragonfly cyclovirus 1 | JX185421.1 |
| Dragonfly cyclovirus 4 | KC512917.1 |
| Dragonfly cyclovirus 6 | KC512918.1 |
| Dhori virus | NC_034255.1 |
| Deformed wing virus, Egypt-1977 isolate | MT504363.1 |
| Deformed wing virus C | CEND01000001.1 |
| Deformed wing virus A, New Zealand isolate | MN538208.1 |
| Varroa destructor virus 1/Deformed wing virus B | AY251269.2 |
| Erysimum latent virus | AF098523.1 |
| Epiphyas postvittana NPV | NC_003083.1 |
| Farmington virus | HM627182.1 |
| Formica exsecta virus 1 | KF500001.1 |
| Formica exsecta virus 2 | KF500002.1 |
| Grapevine Red Globe virus | LC704878.1 |
| Heliconius erato iflavirus | KJ679438.1 |
| Halyomorpha halys virus | KF699344.1 |
| Hubei picorna-like virus 22 | NC_033227.1 |
| Hubei picorna-like virus 29 | KX883292.1 |
| Israel acute paralysis virus of bees | EF219380.1 |
| Jingshan Fly Virus 2 strain JSY-4, partial cds | KM817631.1 |
| Jos virus segment 2 PB1 gene, partial cds | HM627170.1 |
| Kashmir bee virus | AY275710.1 |
| Kakugo virus | AB070959.1 |
| Laodelphax striatellus picorna-like virus 2 | KM272628.1 |
| Lake Sinai virus | KM886903.1 |
| Lake Sinai virus 1 | HQ871931.2 |
| Lake Sinai virus 2 | HQ888865.2 |
| Lake Sinai virus 3 | MH267700.1 |
| Lake Sinai virus 8 | MT482470.1 |
| Mosinovirus strain C36-CI-2004 segment 1 | KJ632942.1 |
| Physalis mottle virus | NC_003634.1 |
| Plantago mottle virus | AY751779.1 |
| Picorna-like virus Eptesicus fuscus, partial genome | HQ585111.1 |
| Slow bee paralysis virus | NC_014137.1 |
| Sacbrood virus | AF092924.1 |
| Scrophularia mottle virus | AY751777.1 |
| Spodoptera exigua Iflavirus-1 | JN091707.1 |
| Thogoto virus | NC_006504.1 |
| Tjuloc virus | JQ928941.1 |
| Tomato blistering mosaic virus | KJ940970.1 |
| Thaumetopoea pityocampa iflavirus 1 | KP217032.1 |
| Turnip yellow mosaic virus | NC_004063.1 |
| Varroa dicistrovirus 1 | MZ822069.1 |
| Apis dicistrovirus 2 | MZ822070.1 |
| Apis dicistrovirus 3 | MZ822071.1 |
| Apis dicistrovirus 4 | MZ822072.1 |
| Varroa destructor Macula-like virus, partial genome | AB859949.1 |
| Varroa destructor virus 2 alpha | this study, PZ380877 |
| Varroa destructor virus 2 beta | this study, PZ380878 |
| Varroa destructor virus 2 delta | this study, PZ380879 |
| Varroa destructor virus 2 epsilon | this study, PZ380880 |
| Varroa destructor virus 2 eta | this study, PZ380881 |
| Varroa destructor virus 2 gamma | this study, PZ380882 |
| Varroa destructor virus 2 theta | this study, PZ380883 |
| Varroa destructor virus 2 zeta | this study, PZ380884 |
| Varroa destructor virus 3 | KX578272.1 |
| Varroa destructor virus 4 | MK032464.1 |
| Varroa destructor virus 5, New Zealand isolate | this study, PZ380885 |
| Varroa destructor virus 9, New Zealand isolate | OR224325.1 |
| Varroa jacobsoni rhabdovirus 1 | MT482464.1 |
| Varroa jacobsoni rhabdovirus 2 | MT482465.1 |
| Varroa jacobsoni virus 2 | MT482466.1 |
| Varroa orthomyxovirus-1 segment, partial genome | OL803864.1 |
| Varroa picorna-like virus | MZ822077.1 |
| Varroa Tymo-like virus | KT162926.1 |
| Wellfeet Bay virus, partial genome | NC_025793.1 |
| Watercress white vein virus | JQ001816.1 |

1. Lester, P.J., Felden, A., Baty, J.W., Bulgarella, M., Haywood, J., Mortensen, A.N., Remnant, E.J., Smeele, Z.E., 2022. Viral communities in the parasite Varroa destructor and in colonies of their honey bee host (Apis mellifera) in New Zealand. Sci Rep 12, 8809. <https://doi.org/10.1038/s41598-022-12888-w>
2. Li, N., Li, C., Hu, T., Li, J., Zhou, H., Ji, J., Wu, J., Kang, W., Holmes, E.C., Shi, W., Xu, S., 2023. Nationwide genomic surveillance reveals the prevalence and evolution of honeybee viruses in China. Microbiome 11, 6. <https://doi.org/10.1186/s40168-022-01446-1>
3. Damayo, J.E., McKee, R.C., Buchmann, G., Norton, A.M., Ashe, A., Remnant, E.J., 2023. Virus replication in the honey bee parasite, Varroa destructor. Journal of Virology 97, e01149-23. <https://doi.org/10.1128/jvi.01149-23>

**Table S3 –** Varroa destructor virus 2 strains assembled from *V. destructor* transcriptomes in New Zealand (NCBI BioProject PRJNA820512), and USA (NCBI BioProject PRJNA380433; SRA accessions SRR5377263-SRR537770).

| **Virus** | **Accession** |
| --- | --- |
| Varroa destructor virus 2 alpha, New Zealand Isolate | this study, PZ380877 |
| Varroa destructor virus 2 beta, New Zealand Isolate | this study, PZ380878 |
| Varroa destructor virus 2 delta, New Zealand Isolate | this study, PZ380879 |
| Varroa destructor virus 2 epsilon, New Zealand Isolate | this study, PZ380880 |
| Varroa destructor virus 2 eta, New Zealand Isolate | this study, PZ380881 |
| Varroa destructor virus 2 gamma, New Zealand Isolate | this study, PZ380882 |
| Varroa destructor virus 2 theta, New Zealand Isolate | this study, PZ380883 |
| Varroa destructor virus 2 zeta, New Zealand Isolate | this study, PZ380884 |
| Varroa destructor virus 2 alpha, USA Isolate | this study, BK083176 |
| Varroa destructor virus 2 beta, USA Isolate | this study, BK083173 |
| Varroa destructor virus 2 delta, USA Isolate | this study, BK083180 |
| Varroa destructor virus 2 epsilon, USA Isolate | this study, BK083175 |
| Varroa destructor virus 2 eta, USA Isolate | this study, BK083174 |
| Varroa destructor virus 2 gamma, USA Isolate | this study, BK083177 |
| Varroa destructor virus 2 theta, USA Isolate | this study, BK083179 |
| Varroa destructor virus 2 zeta, USA Isolate | this study, BK083178 |

**Table S4**. Primer sequences used for qPCR. Primers were designed using Primer3 in Geneious Prime. 18S and NADHD were used as reference genes as their stability has been previously validated in all *V. destructor* life stages^1^. All *V. destructor* primers span exon-exon junctions to ensure specificity to mRNA targets. Primer efficiencies were determined by five, 3-fold serial dilutions

| Target Sequence | Sequence  (5 to’3’) | Primer concentration  (nM) | Primer efficiency | Product size (bp) |
| --- | --- | --- | --- | --- |
| 18S  (LOC111253957) | **F:** CAAACGAAGGTCATGTATGCC | 300 | 105.28% | 132 |
|  | **R:** GGCATTCACTTCCTGTTCCG | 300 |  |  |
| NADHD (LOC111249888) | **F:** CGAGTTCTATAAGCACGAGAGC | 200 | 100.20% | 158 |
|  | **R:** TGGTATGACCCTCAATCTGC | 200 |  |  |
| Ago-2 (LOC111247833) | **F:** CGGGATCAACACAGAGTTCG | 200 | 127.12% | 124 |
|  | **R:** AATCGTGGATGGTAGCTTCG | 200 |  |  |
| DWV-A^2^ | **F:** TACTAGTGCTGGTTTTCCTTT | 125 | 97.95% | 155 |
|  | **R:** CTCATTAACTGTGTCGTTGAT | 125 |  |  |

1. Campbell, E.M., McIntosh, C.H., Bowman, A.S., 2016. A Toolbox for Quantitative Gene Expression in Varroa destructor: RNA Degradation in Field Samples and Systematic Analysis of Reference Gene Stability. PLOS ONE 11, e0155640. <https://doi.org/10.1371/journal.pone.0155640>
2. Kevill, J.L., Highfield, A., Mordecai, G.J., Martin, S.J., Schroeder, D.C., 2017. ABC Assay: Method Development and Application to Quantify the Role of Three DWV Master Variants in Overwinter Colony Losses of European Honey Bees. Viruses 9, 314. <https://doi.org/10.3390/v9110314>

**Table S5**. Double stranded RNA sequences used for dsRNA immersion assays in *V. destructor* foundresses. Sequences were designed to span an exon-exon junction within the coding sequence. All constructs were synthesised from in vitro transcription and magnetic bead purification by RNA Greentech LLC (Texas, USA).

| Target Sequence | Sequence  (5 to’3’) | Product size (bp) |
| --- | --- | --- |
| Ago-2 (LOC111247833) | GTGTTGCGAGAAATGACGTTCCCACAGCTCTGCATAGAAAACAAATTTTAGAACTGTCCAAAAAGCTTTCACGGTGCAAAGCAGAGCTTACGCATCTCAAAAGACCTAAACATATTACCATACAAAAAATTACCCTTGAGCCTGCAAGCAAAATTTCATTCAGAGATCATGAGAACAACCCTATCCTTGTGACAGATTACTTCAGCAAAACCTATGGTTGCTTGAAATATCCAAACTGGCCTTGTGCGGAGGTCGGTGTTCGAGAACCTAGATATTACCCGCTAGAGGTTTTGAAAATACTGAGCGGTAATTCCTACCACGGAGAACAATCACGTGATATGGTGCGCGACATAATACGTTTGGCGGCCATTCCTCCGCGCAATAGGCTGGAAGAAATTAAATTCACCACGCGCTCGTTGATAAGTCACAAAGAGATAAAGGATGAGTTCGGTATGCATATTGAACCTATCGAAATGACGATTCCTGGAAGAATTTTGCCCACACCGATACTTACAGCAGGAAACAAGAAAATTGTCGGAGTTAACCCTGGGGTTATCGATTTCAGAGACAGTATGTTCTTTAAAACATCCAAGATTGATCGGTGGACAGTAATTAACACCGATGAAAGGCAGACACAGGATACGTTGCAGTGCTTTGCCAGAGCATTTCAGAACCAAGGAAAAAAAAGCGGCGTACTGCTTGCAAATCCTGTGCAGCACAACAACGAATTTCCAACCATAAGCGACCGTCAGGTGCTGAAAAACATCATACAACAGATCATTAAGATGGTTACTGTGAAC | 800 |
| GFP (L29345.1) | GGGCAAAAATTCTCTGTCAGTGGAGAGGGTGAAGGTGATGCAACATACGGAAAACTTACCCTTAAATTTATTTGCACTACTGGGAAGCTACCTGTTCCATGGCCAACACTTGTCACTACTTTCTCTTATGGTGTTCAATGCTTTTCAAGATACCCAGATCATATGAAACAGCATGACTTTTTCAAGAGTGCCATGCCCGAAGGTTATGTACAGGAAAGAACTATATTTTACAAAGATGACGGGAACTACAAGACACGTGCTGAAGTCAAGTTTGAAGGTGATACCCTTGTTAATAGAATCGAGTTAAAAGGTATTGATTTTAAAGAAGATGGAAACATTCTTGGACACAAAATGGAATACAACTATAACTCACATAATGTATACATCATGGCAGACAAACCAAAGAATGGAATCAAAGTTAACTTCAAAATTAGACACAACATTAAAGATGGAAGCGTTCAATTAGCAGACCATTATCAACAAAATACTCCAATTGGCGATGGCCCTGTCCTTTTACCAGACAACCATTACCTGTCCACACAATCTGCCCTTTCCAAAGATCCCAACGAAAAGAGAGATCACATGATCCTTCTTGAGTTTGTAACAGCTGCTGGGATTACACATGGCATGGATGAACTATACAAATAAATGTCCAGACTTCCAATTGACACTAAAGTGTCCGAACAATTACTAAATTCTCAGGGTTCCTGGTTAAATTCAGGCTGAGACTTTATTTATATATTTATAGATTCATTAAAATTTTATGAATAATTTATTGATGTTATTAATAGGGGCTATTT | 800 |

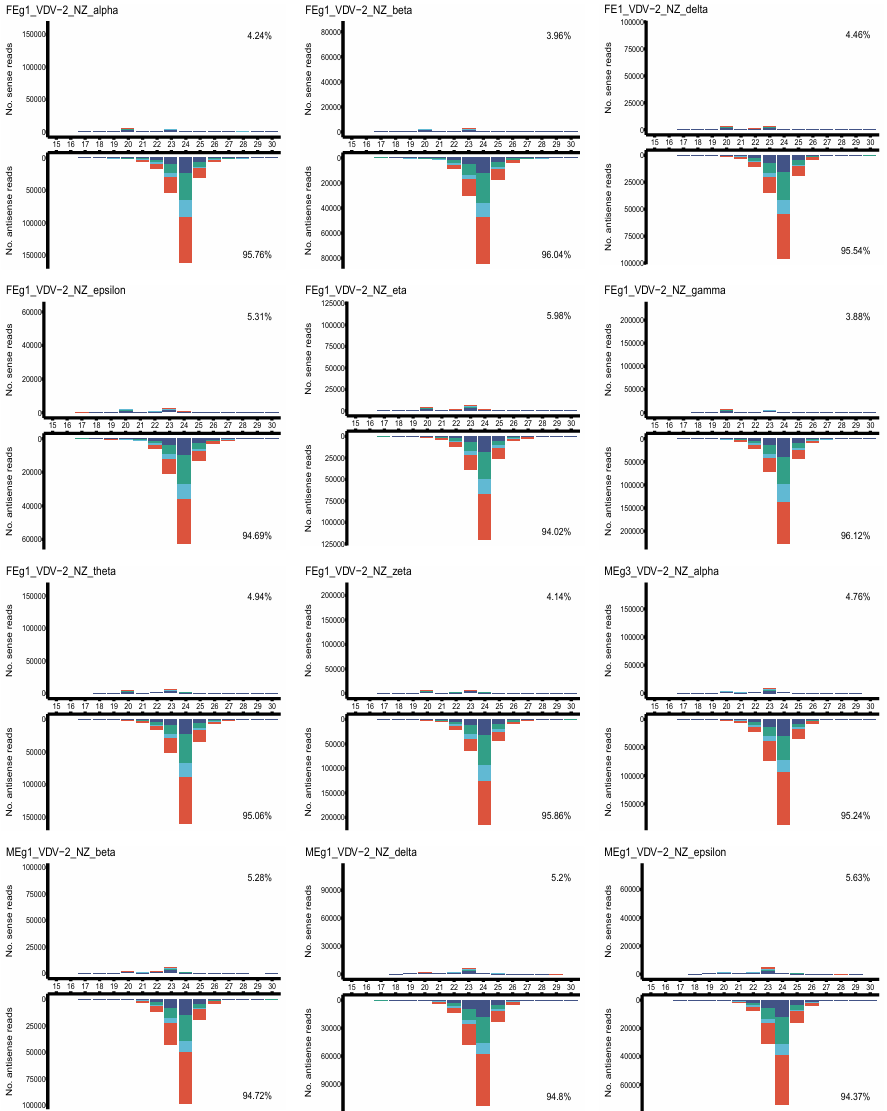
**Figure S1** continued to the next page; legend on page 9

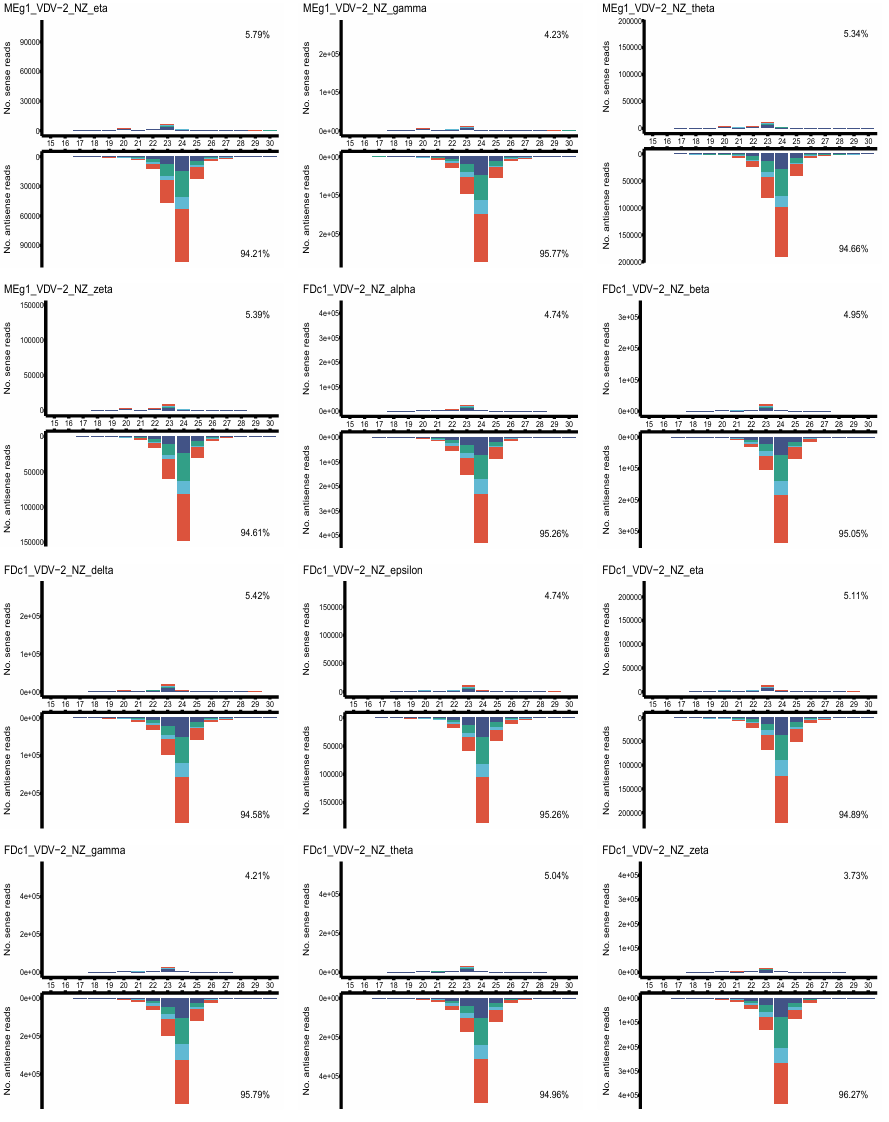
 **Figure S1** continued to next page; legend on page 9

**Figure S1** continued on next page; legend on page 9
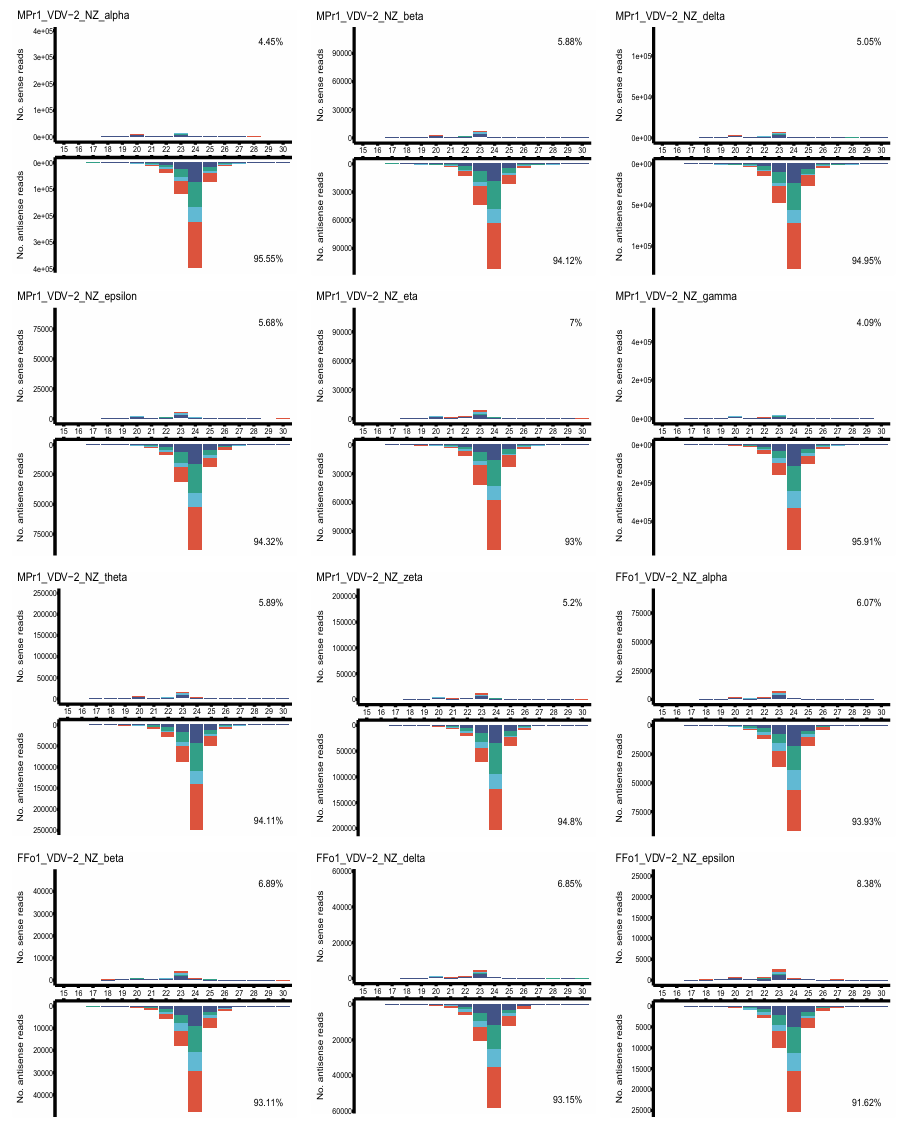

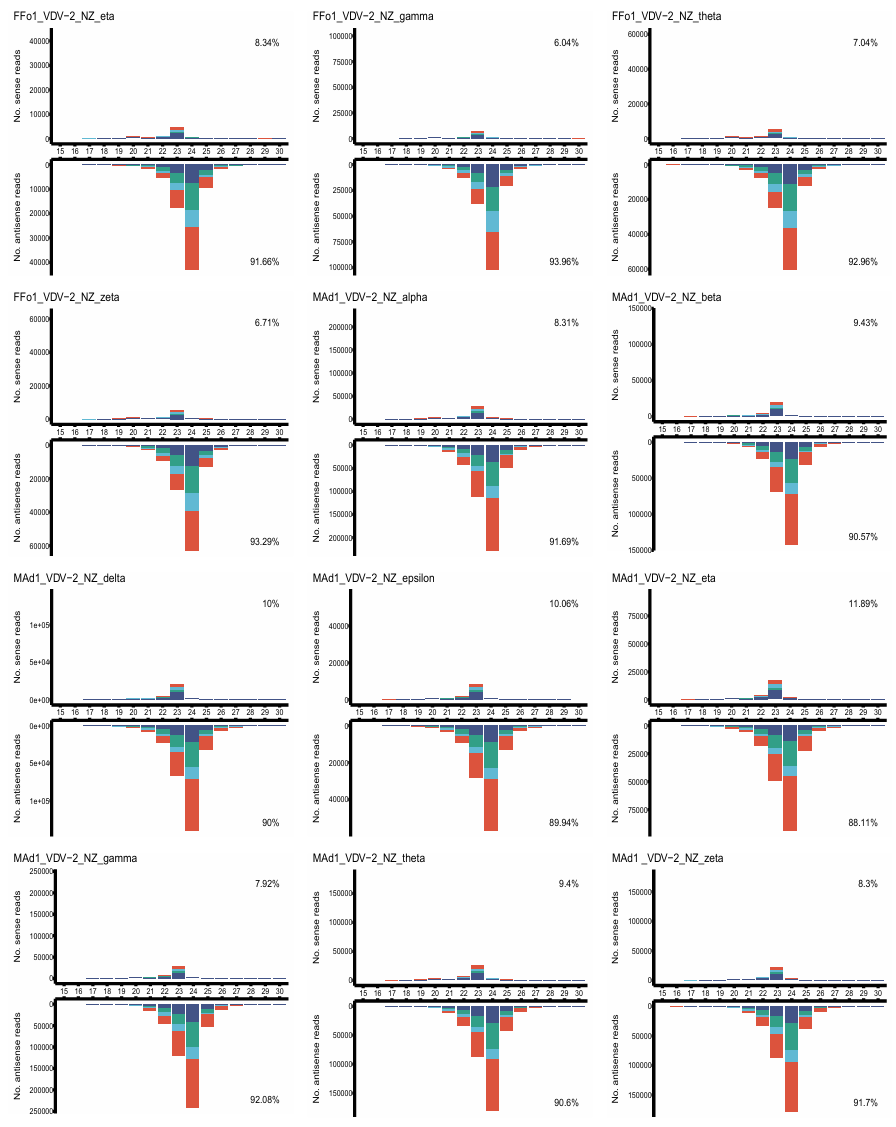

**Figure S1** – Virus derived siRNA profiles of Varroa destructor virus 2 (VDV-2) strains: -alpha, -beta, -delta, -epsilon, -eta, -gamma, -theta, and -zeta strains. All VDV-2 strains were present in all samples. Profiles generated from single, representative samples from a female egg (FEg1) and male egg (MEg1), female deutonymph (FDc1), male protonymph (MPr1), female adult (FAd1), and male adult (MAd1).

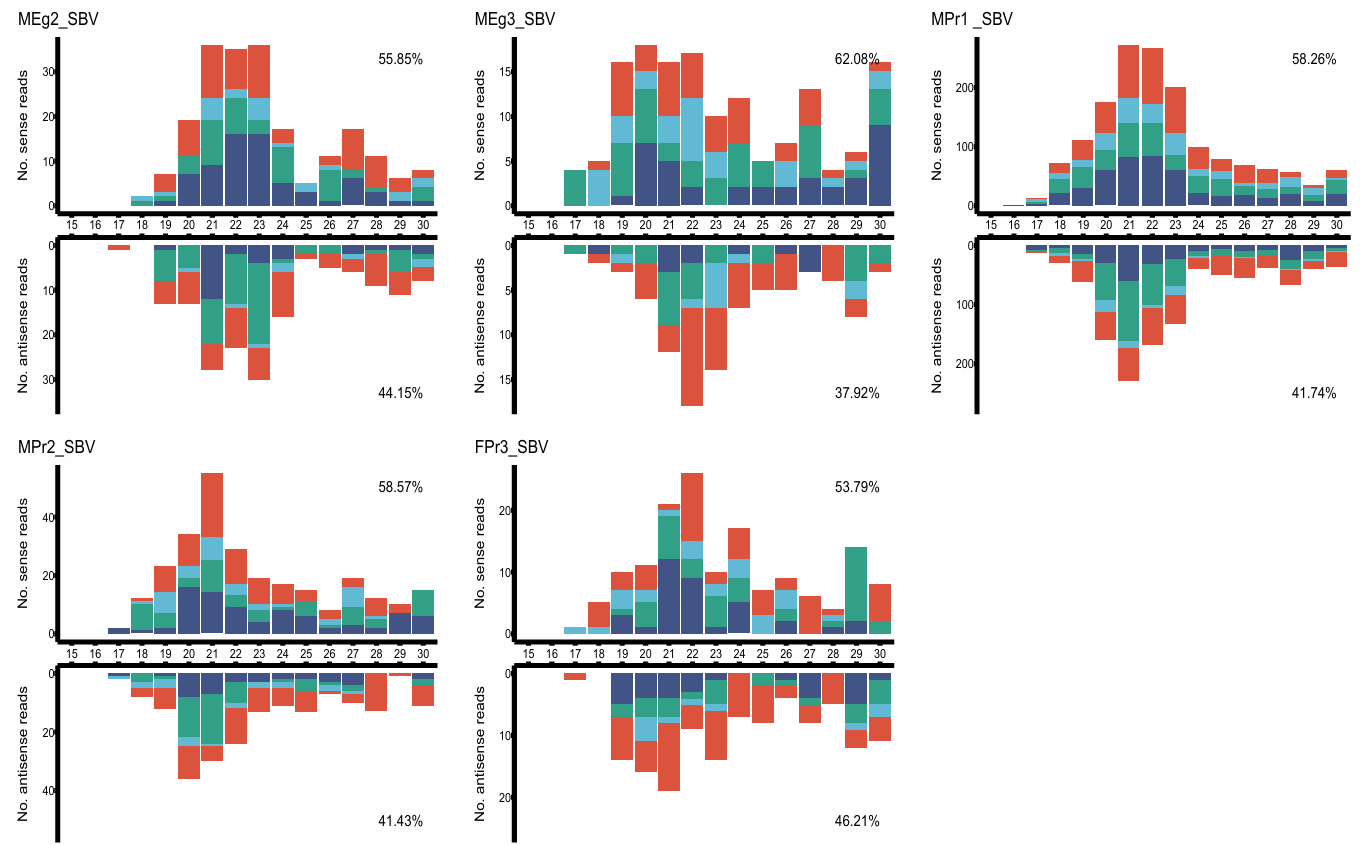

**Figure S2** – Virus derived siRNA profiles of sacbrood virus (SBV, AF092924.1) in *V. destructor* samples (MEg2, MEg3, MPr1, MPr2, and FPr3). Profiles were presented if at least 100 reads were mapped, had greater than 30% identity, and exceeded 1 RPM (reads per million).

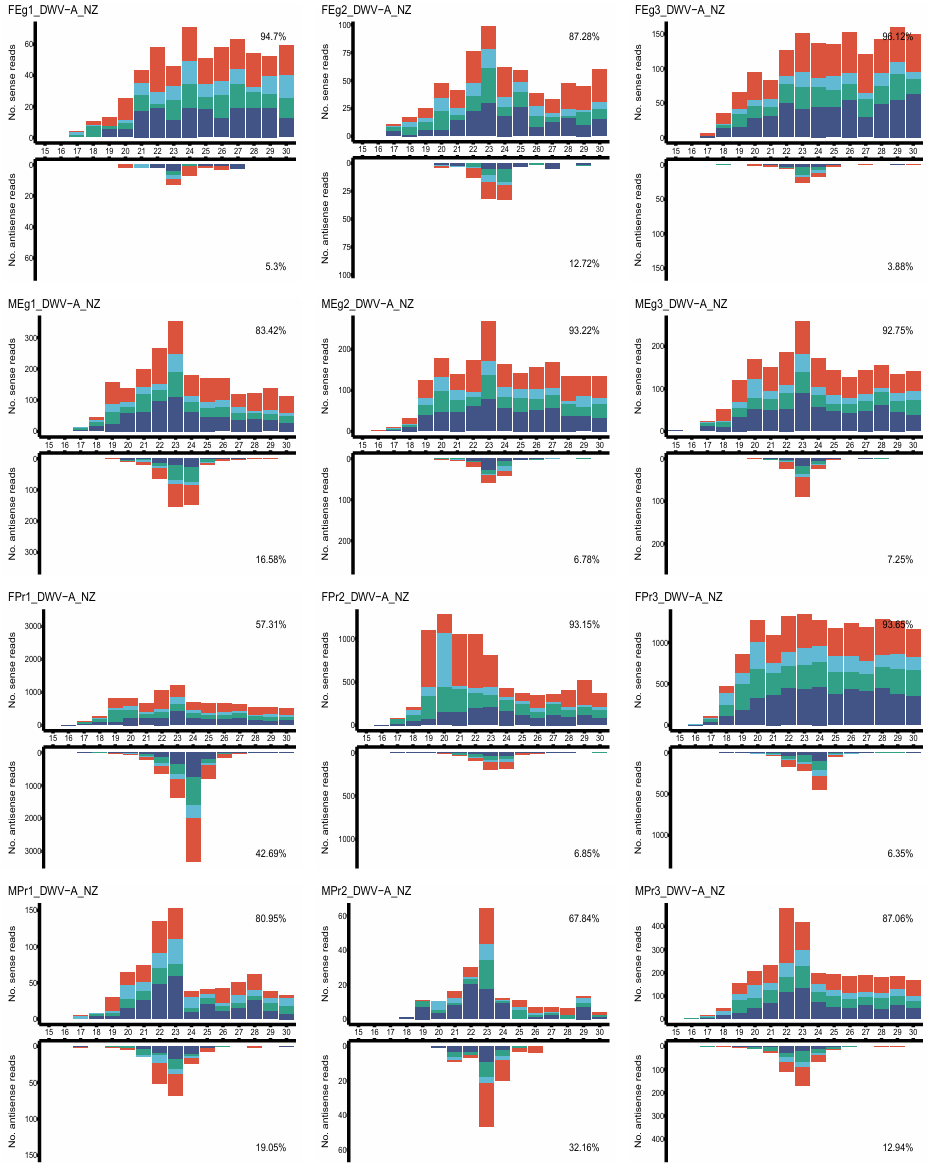

**Figure S3** continued on next page; legend on page 14

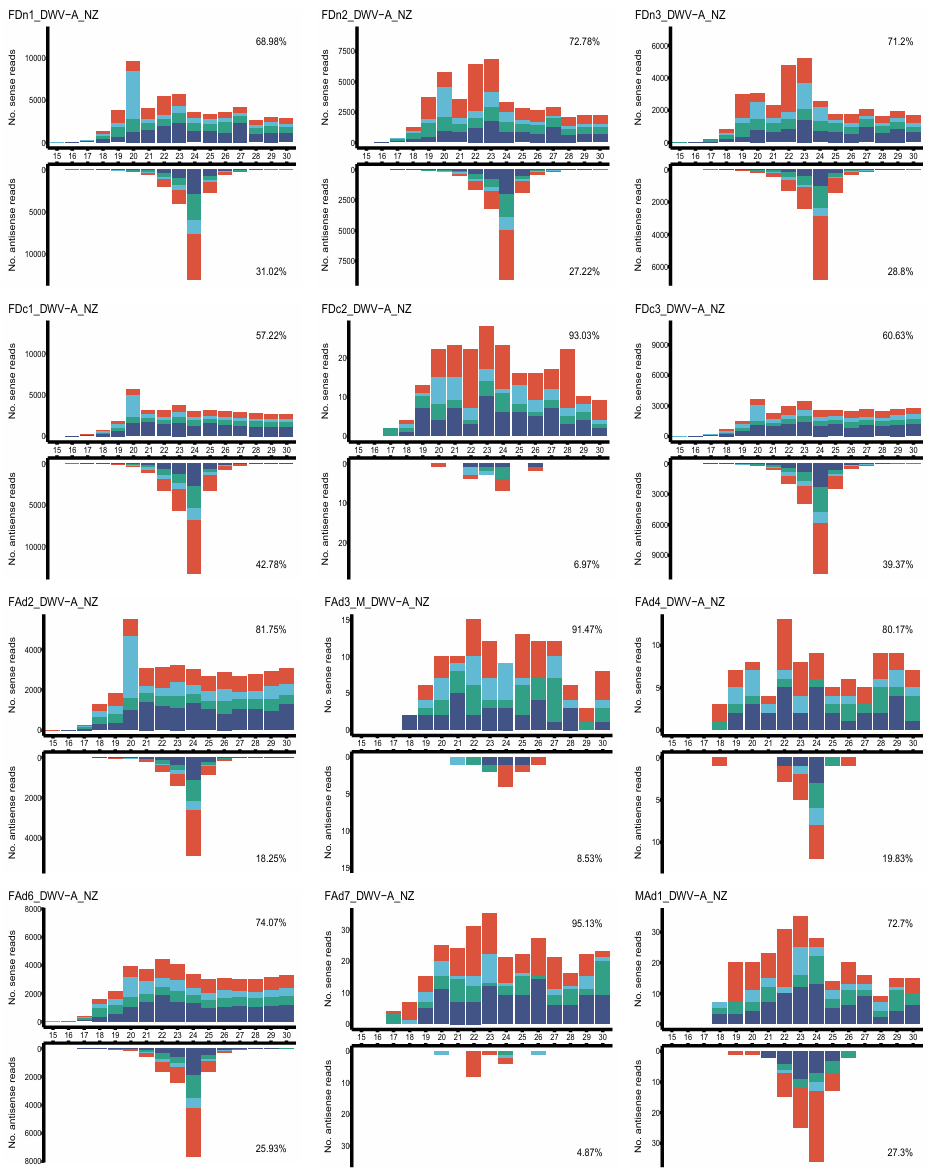

**Figure S3** continued next page; legend on page 14

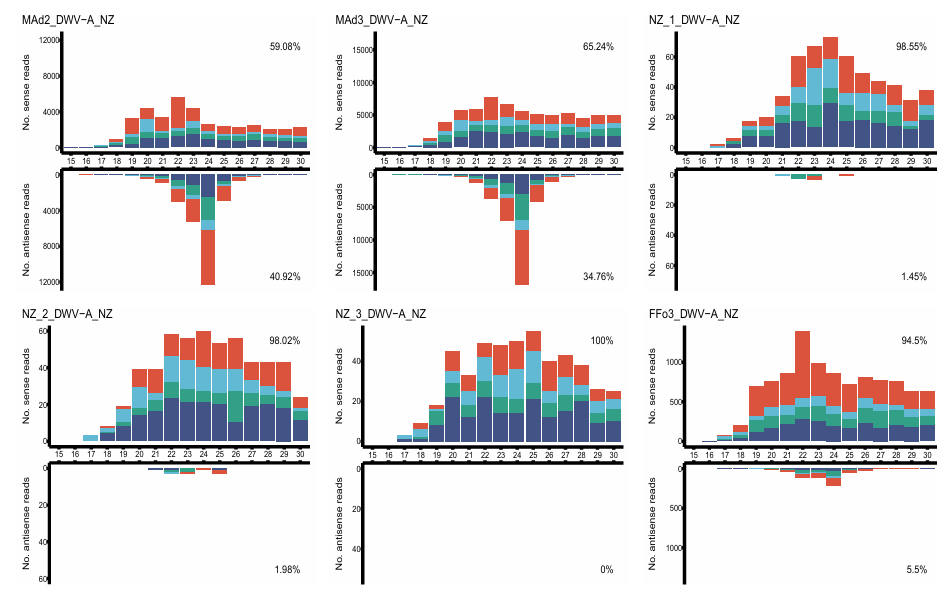

**Figure S3** – Virus derived siRNA profiles of deformed wing virus A (DWV-A), New Zealand isolate (MN538208.1)in *V. destructor* life stage samples. Life stages include female eggs (FE), male eggs (ME), female protonymph (FPr), male protonymph (MPr), female deutonymph (FDn), female pre-deutochrysalis (FDc), young female adult (FAd), female foundress (FFo), and dispersal female (FDi). Profiles were presented if at least 100 reads were mapped, had greater than 30% identity, and exceeded 1 RPM (reads per million).

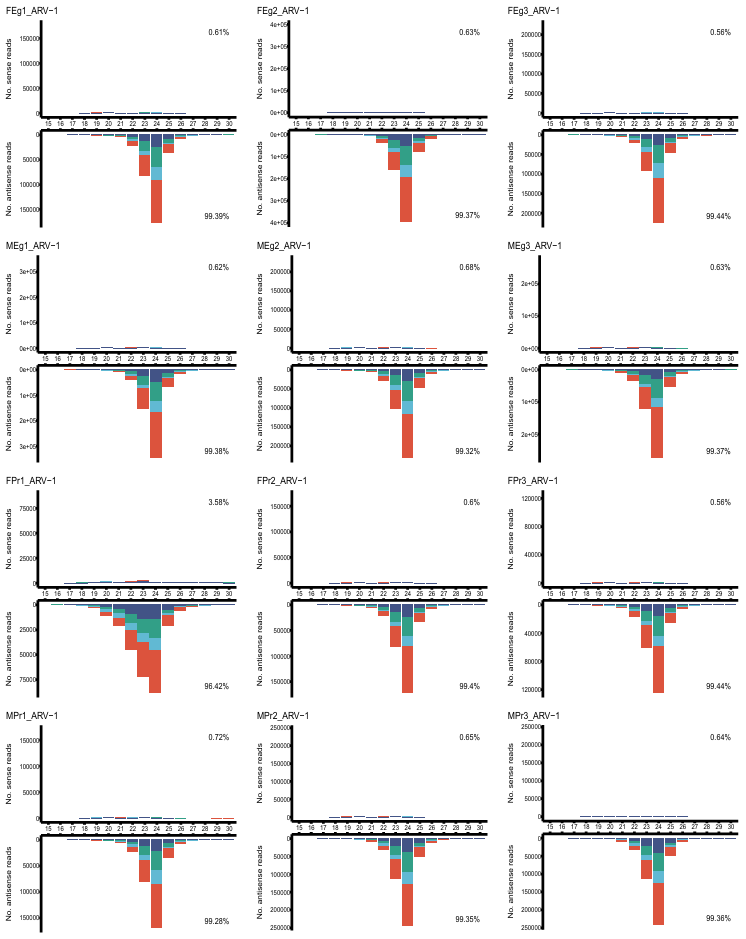
**Figure S4** continued next page; legend on page 17

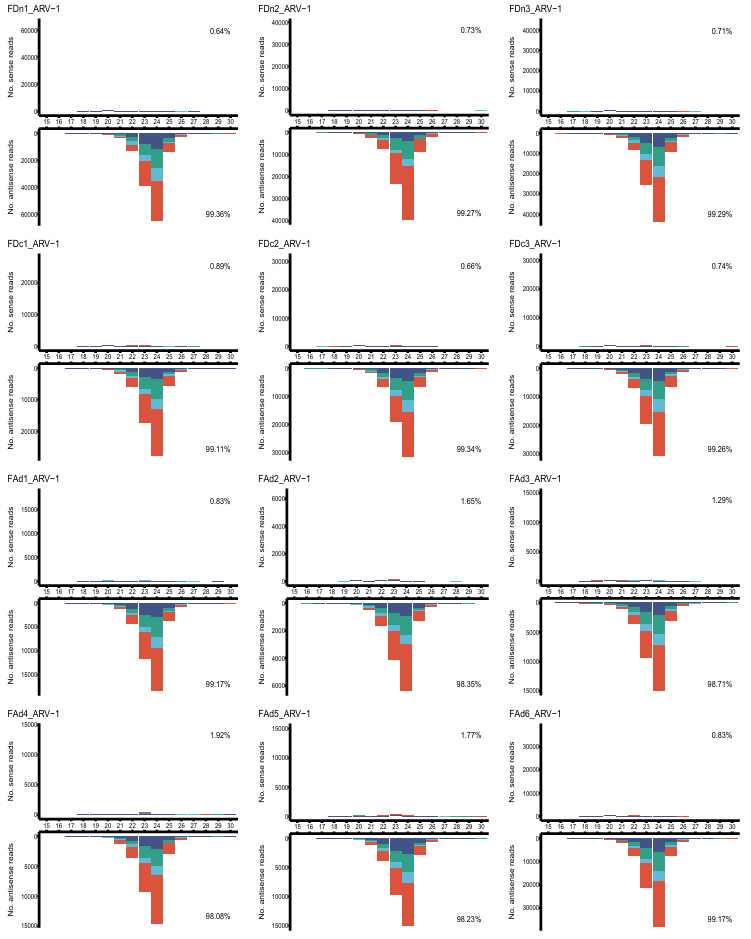
**Figure S4** continued next page; legend on page 17

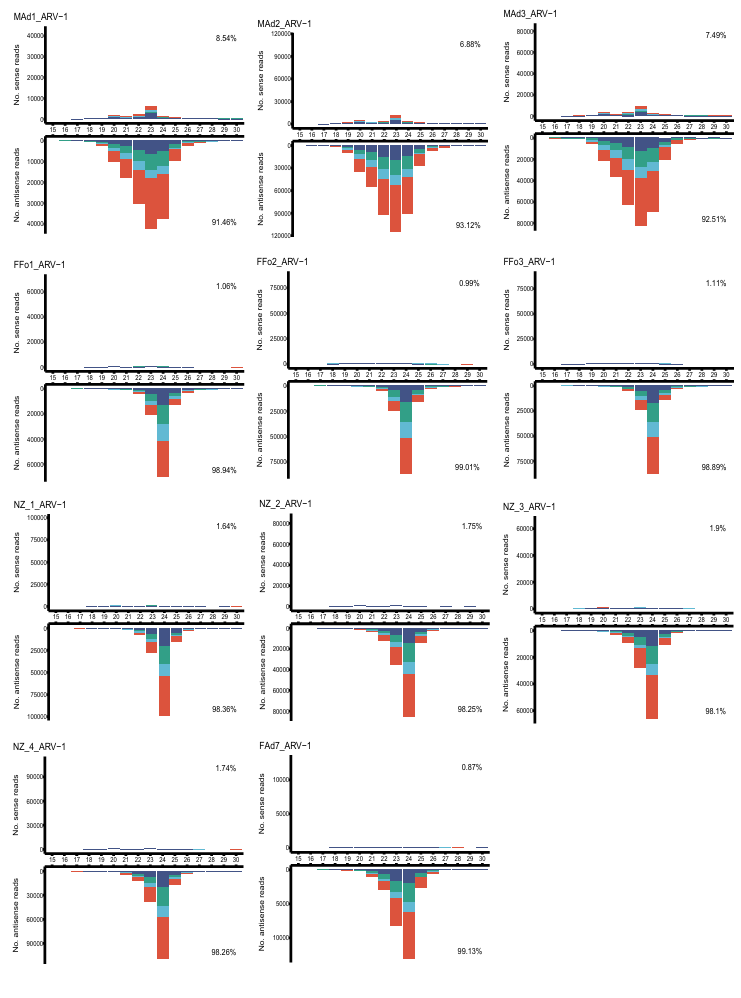
**Figure S4** – vsiRNA profiles of Apis rhabdovirus 1 (ARV-1, MF114351.1) in *V. destructor* life stage samples. Life stages include female eggs (FE), male eggs (ME), female protonymph (FPr), male protonymph (MPr), female deutonymph (FDn), female pre-deutochrysalis (FDc), young female adult (FAd), female foundress (FFo), and dispersal female (FDi). Profiles were presented if at least 100 reads were mapped, had greater than 30% identity, and exceeded 1 RPM (reads per million).

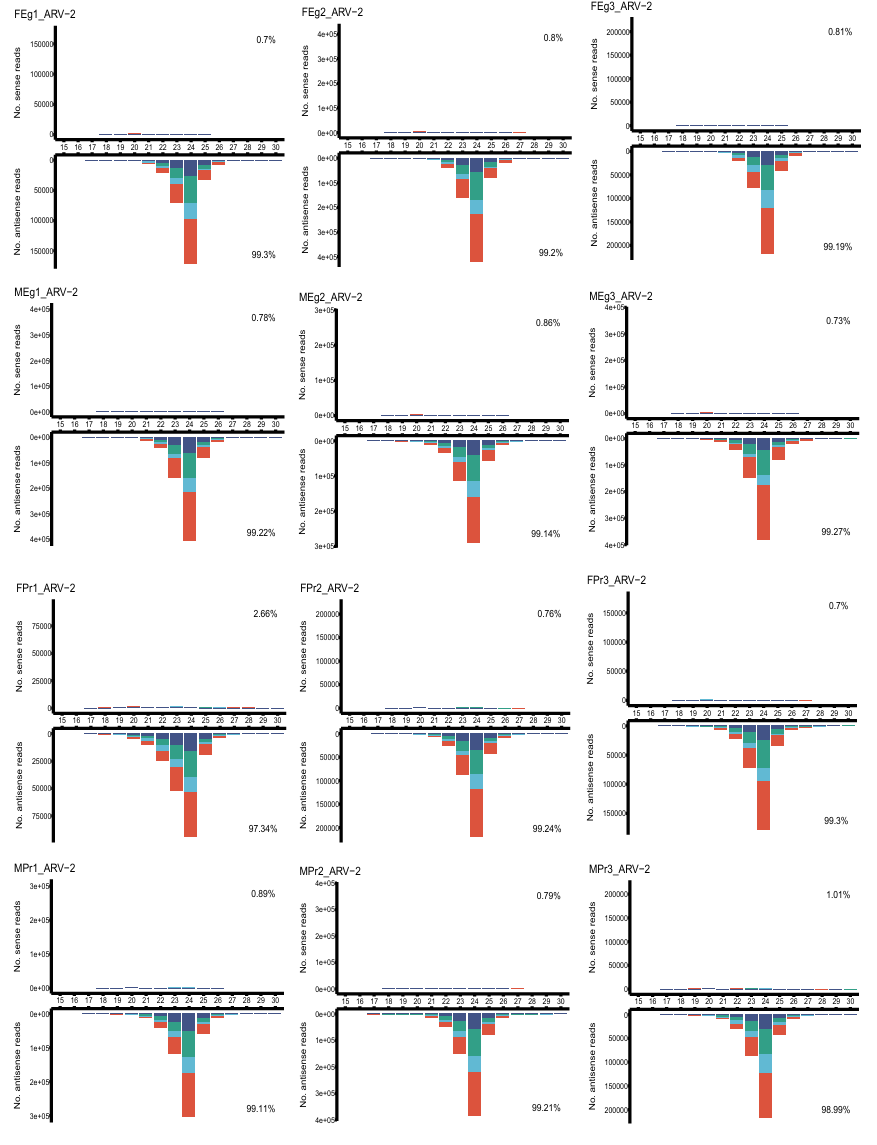
**Figure S5** continued next page; legend on page 20

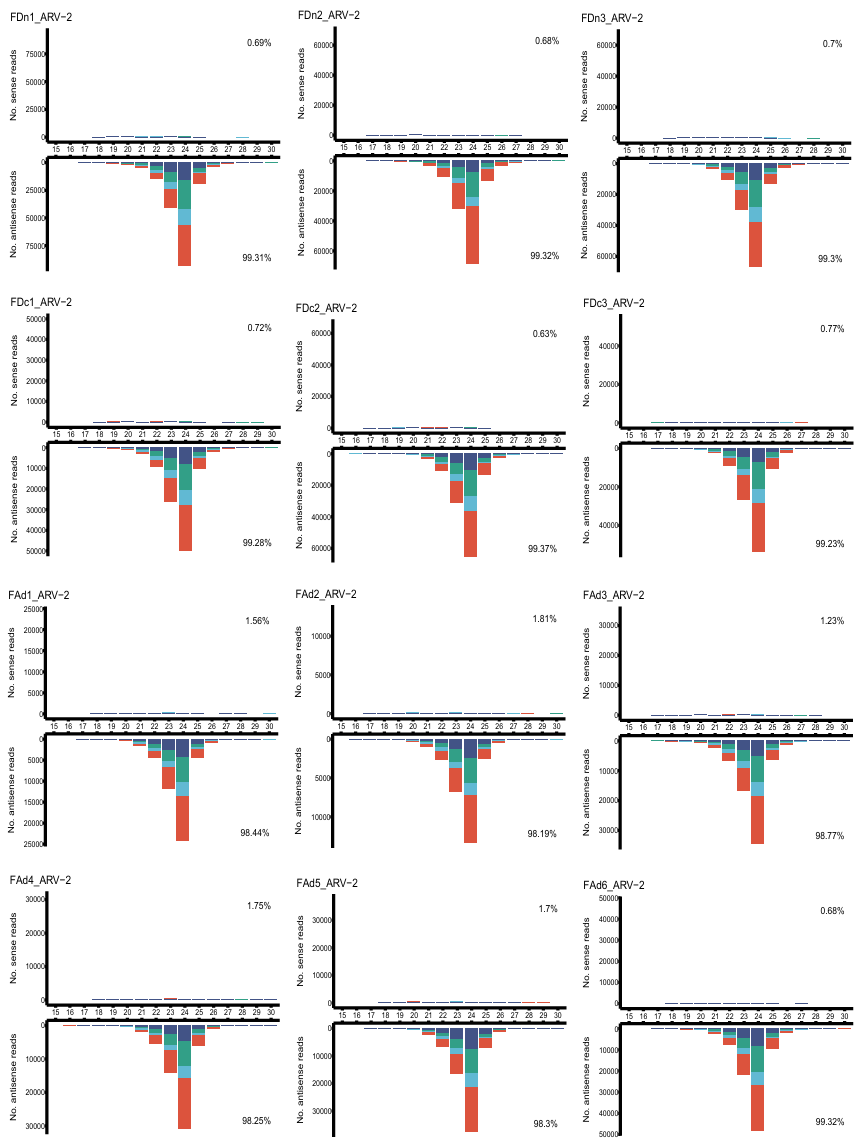
**Figure S5** continued next page; legend on page 20

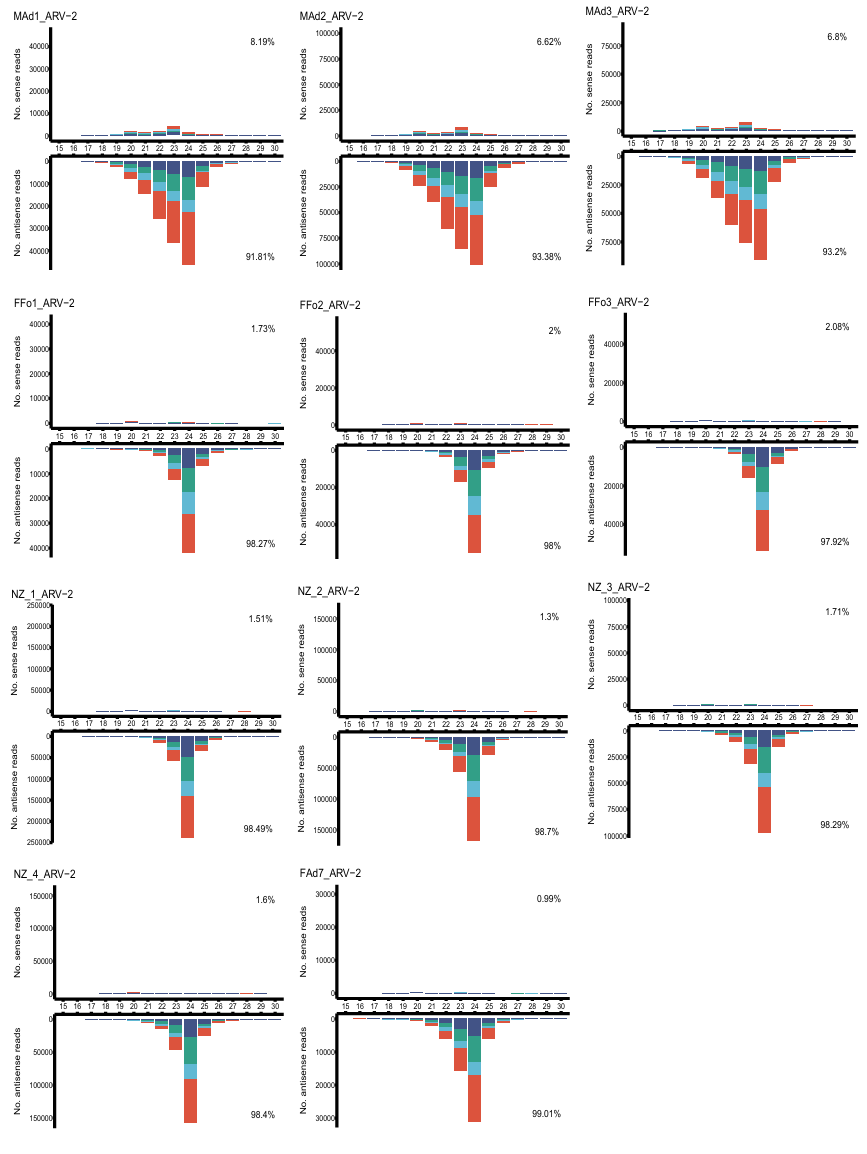

**Figure S5** – Virus derived siRNA profiles of Apis rhabdovirus 2 (ARV-2, MZ821796.1) in *V. destructor* life stage samples. Life stages include female eggs (FE), male eggs (ME), female protonymph (FPr), male protonymph (MPr), female deutonymph (FDn), female pre-deutochrysalis (FDc), young female adult (FAd), female foundress (FFo), and dispersal female (FDi). Profiles were presented if at least 100 reads were mapped, had greater than 30% identity, and exceeded 1 RPM (reads per million).

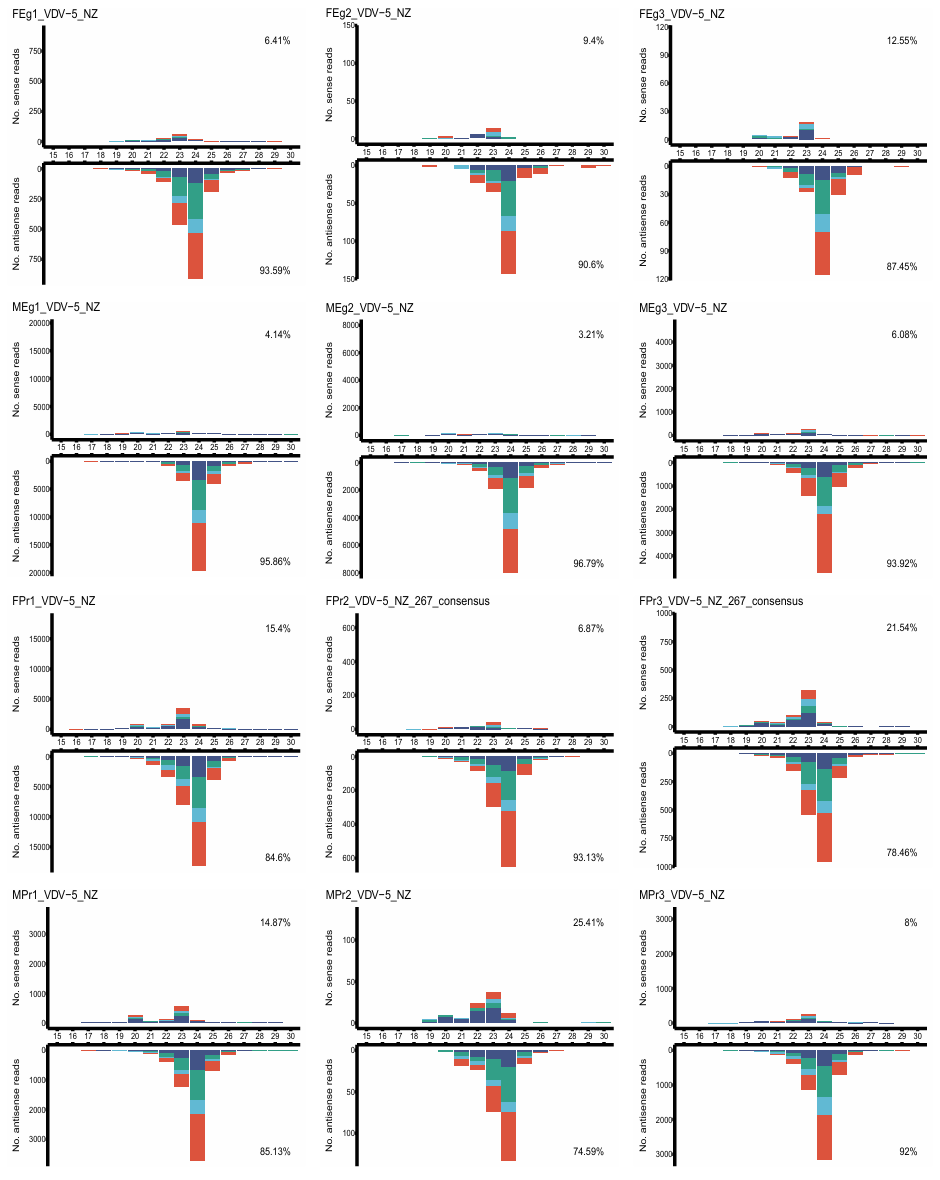
**Figure S6** continued next page; legend on page 23

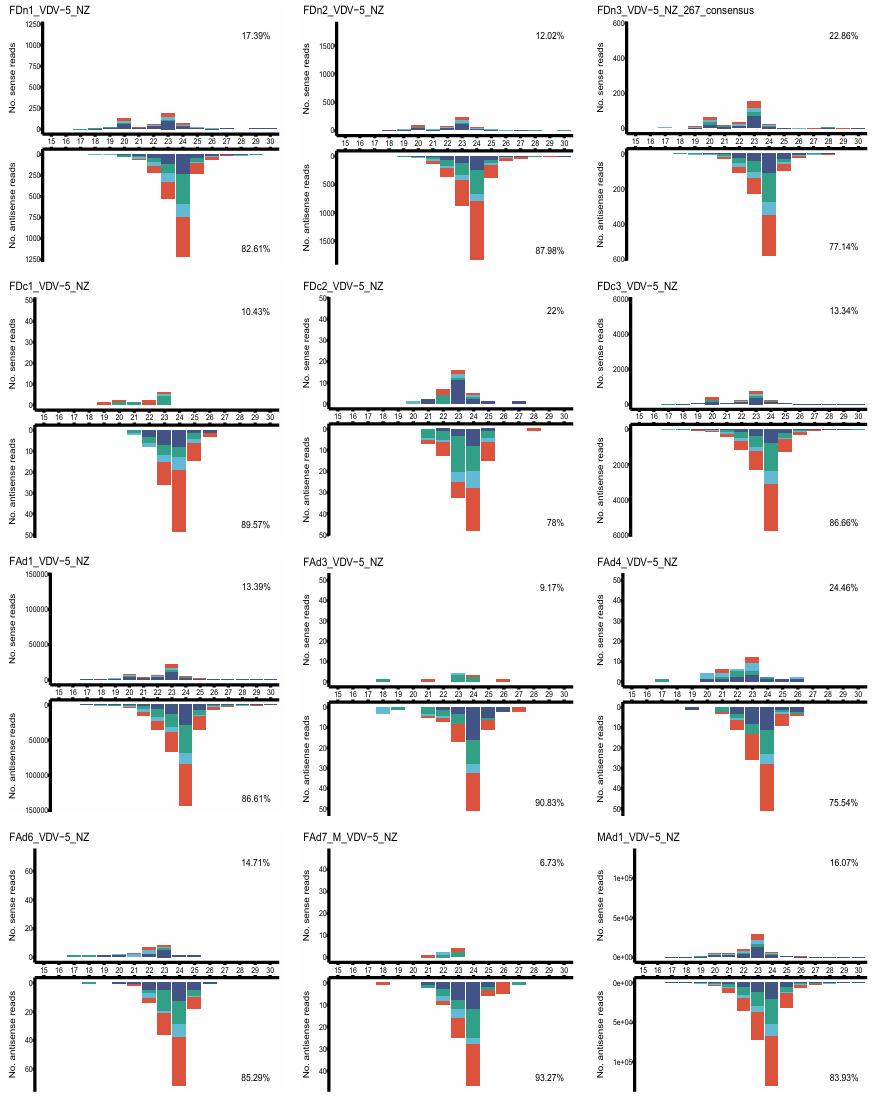
**Figure S6** continued next page; legend on page 23

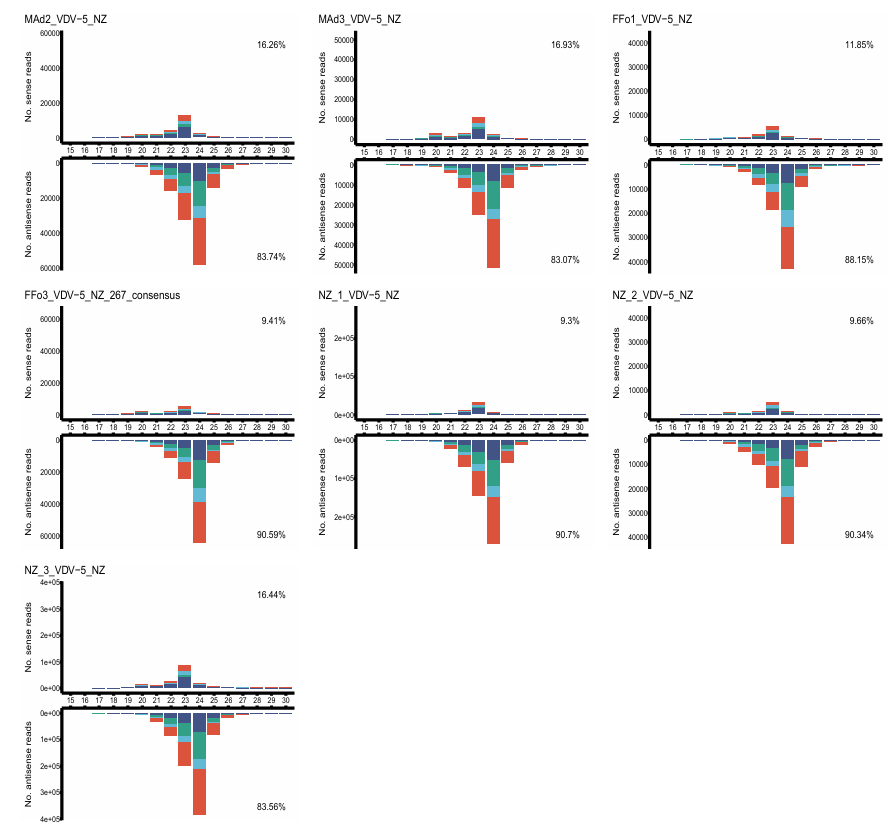

**Figure S6** – Virus derived siRNA profiles of Varroa destructor virus 5, New Zealand isolate (VDV-5_NZ, accession?) isolate in *V. destructor* life stage samples. Life stages include female eggs (FE), male eggs (ME), female protonymph (FPr), male protonymph (MPr), female deutonymph (FDn), female pre-deutochrysalis (FDc), young female adult (FAd), female foundress (FFo), and dispersal female (FDi). Profiles were presented if at least 100 reads were mapped, had greater than 30% identity, and exceeded 1 RPM (reads per million).

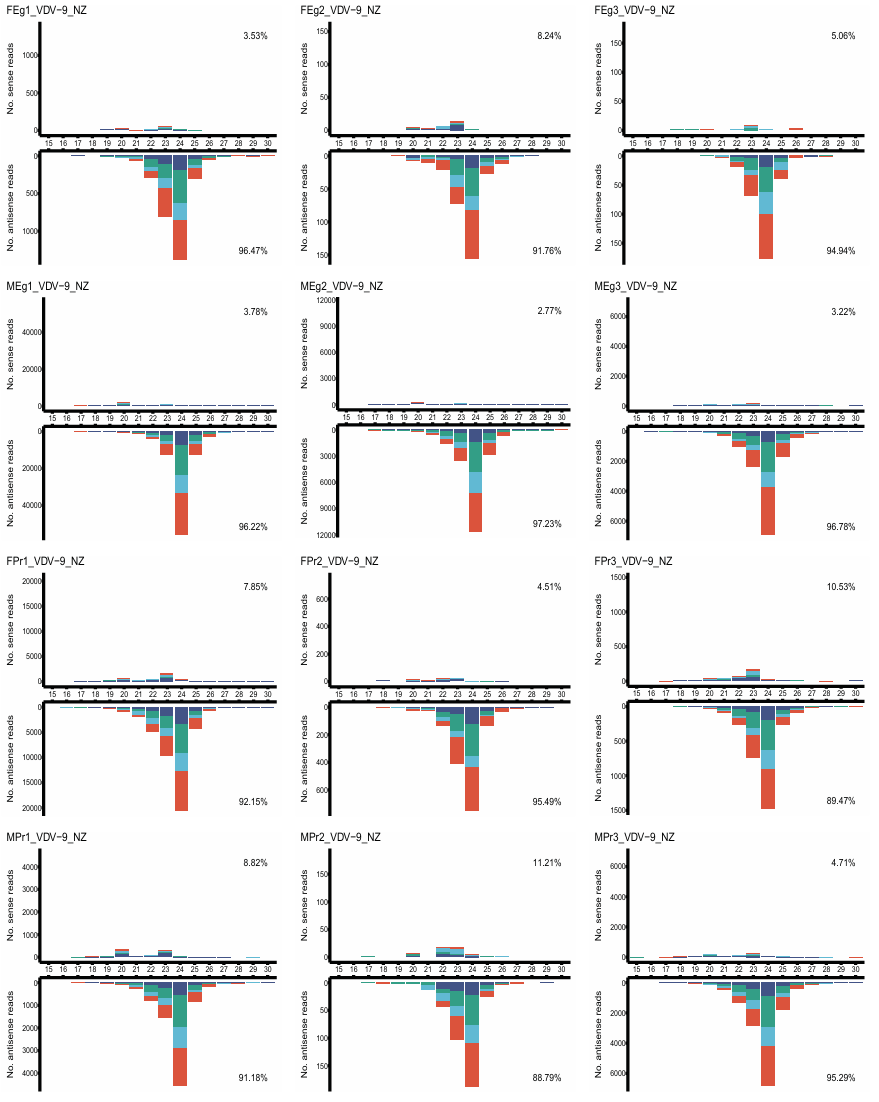
**Figure S7** continued next page; legend on page 26

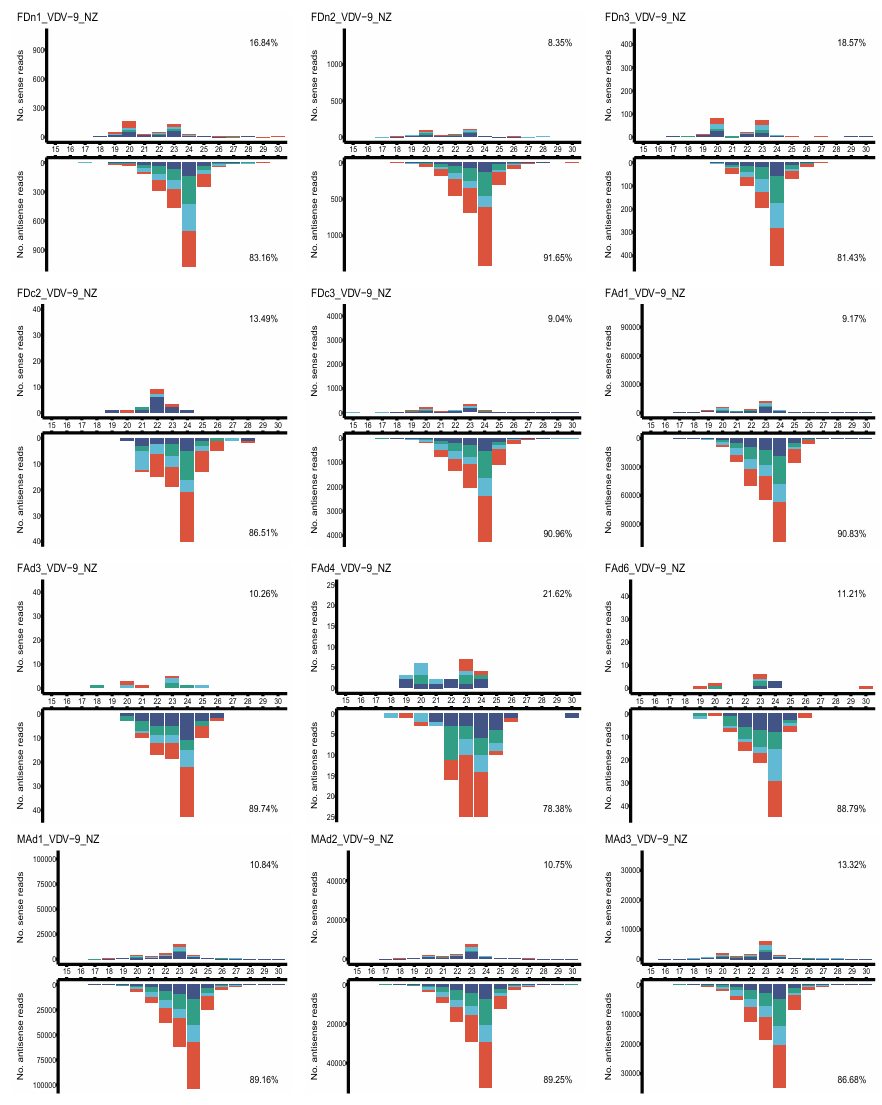
**Figure S7** continued next page; legend on page 26

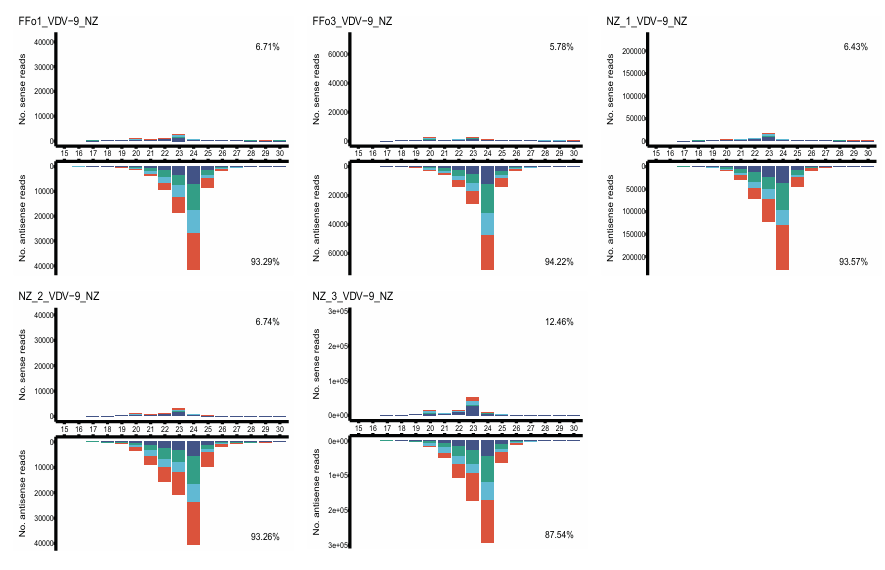

**Figure S7** – Virus derived siRNA profiles of Varroa destructor virus 9, New Zealand isolate (VDV-9_NZ, OR224325.1) isolate in *V. destructor* life stage samples. Life stages include female eggs (FE), male eggs (ME), female protonymph (FPr), male protonymph (MPr), female deutonymph (FDn), female pre-deutochrysalis (FDc), young female adult (FAd), female foundress (FFo), and dispersal female (FDi). Profiles were presented if at least 100 reads were mapped, had greater than 30% identity, and exceeded 1 RPM (reads per million).
